# An endoplasmic reticulum (ER) ATPase safeguards ER identity by removing ectopically localized mitochondrial proteins

**DOI:** 10.1101/2020.09.27.316026

**Authors:** Qing Qin, Ting Zhao, Wei Zou, Kang Shen, Xiangming Wang

## Abstract

Stringent targeting of membrane proteins to corresponding organelles is essential for organelle identity and functions. In addition to molecular pathways that target proteins to appropriate organelles, surveillance mechanisms clear mistargeted proteins from undesired destinations. While Msp1 functions on mitochondrial membrane to remove mistargeted proteins, the surveillance mechanism for the ER is not well understood. Here, we show that mitochondrial tail-anchored (TA) and signal-anchored (SA) proteins mislocalize to ER membrane in neurons and muscles in *C. elegans catp-8* mutants. *catp-8* encodes a conserved P5A type ATPase, which localizes to ER and removes ectopic mitochondrial TA/SA proteins from ER. In *catp-8* mutant, mitochondria fission protein FIS-1 mislocalizes to ER membrane. Together with another mitochondria fission protein MFF-2, FIS-1 causes ER fragmentation in a Dynamin related protein (DRP-1) dependent manner. Additionally, CATP-8 is essential for dendrite development. *catp-8* mutant dramatically reduces the level of the dendrite guidance receptor DMA-1, leading to diminished dendritic arbors. Hence, P5A ATPase safeguards ER morphology and functions by preventing mitochondrial proteins mislocalization.

**HIGHLIGHTS:** CATP-8, a P5A type ATPase, localizes to ER and functions as a surveillance mechanism to remove mistargeted mitochondrial proteins.

Multiple mitochondria proteins are mistargeted to ER in *catp-8* mutants.

Ectopic recruitment of mitochondria fission mechinary to ER causes ER fragmentation in *catp-8* mutants.

CATP-8 is essential for PVD dendrite morphogenesis through modulating the level of transmembrane receptor DMA-1.

## INTRODUCTION

Targeting proteins to the appropriate organelles is essential for cellular organization and organelle function. Signal recognition particle (SRP) guided co-translational translocation across the ER membrane represents the major way for proteins to be inserted into lipid bilayers (Nyathi et al., 2013). Post-translational insertion mechanisms are also important for certain classes of transmembrane proteins, especially for ER or mitochondrial proteins (Chacinska et al., 2009; Costa et al., 2018; Johnson et al., 2013).

One class of such proteins is the tail-anchored (TA) proteins, which are inserted to the ER or mitochondria membrane through a single transmembrane domain at the extreme C-terminus of their coding sequences. Several mechanisms have been reported to target TA proteins to ER. The GET (guided entry of TA proteins) pathway is the dominant cellular mechanism for inserting newly synthesized TA proteins to ER (Wang et al., 2011). In budding yeast, Sgt2, a TA protein chaperone forms a complex with Get4 and Get5 to recognize the C-terminal transmembrane domain (TMD) signals for ER targeting (Chang et al., 2010). Subsequently, the TA proteins are transferred from Sgt2 to Get3, an ATPase chaperone which hands the TA proteins over to the Get1/2 complex for ER membrane insertion (Wang et al., 2014). A recent study shows that the ER membrane protein complex (EMC) also functions as an ER transmembrane domain insertase (Guna et al., 2018). The mitochondria outer membrane (OMM) also contains many important TA proteins, however the targeting mechanism for mitochondrial TA proteins remains largely unknown. It is also not well understood what underlies the selective targeting of TA proteins to ER or mitochondria. One hypothesis is that the low ergosterol concentration of OMM makes it possible for TA proteins to be inserted spontaneously without any specific insertion mechanims (Brambillasca et al., 2005; Kemper et al., 2008; Krumpe et al., 2012). This is supported by the fact that impairment of the GET pathway leads to the mistargeting of a subset of ER-destined TA proteins to mitochondria (Okreglak and Walter, 2014).

In the OMM, another class of transmembrane proteins is anchored in the membrane through an N-terminal α-helix, which functions as the targeting signal with the flanking positively charged amino acid residues (Wasilewski et al., 2017; Wiedemann and Pfanner, 2017). These proteins are collectively called signal-anchored (SA) proteins and include proteins such as Tom20 and Tom70, which expose the C-terminal sequences to the cytosol and function as mitochondrial protein import machines for inner membrane and matrix proteins (Chacinska et al., 2009). In yeast, the mitochondrial import (MIM) complex, which consists of Mim1 and Mim2, is responsible for targeting precursors of SA proteins to OMM (Becker et al., 2008). A recent paper reports that MIM also facilitates the membrane insertion of the TA protein Gem1 (Vitali et al., 2020). However, no homologs of the MIM complex are found in metazoans. No systematic understanding of the targeting mechanisms for SA proteins exists for metazoans (Wasilewski et al., 2017).

The fidelity of targeting TA/SA proteins to mitochondria and ER is critical for the normal function of these organelles. In addition to the specific ER targeting mechanisms, a surveillance mechanism for mistargeted TA proteins to OMM has been reported. The conserved AAA-ATPase Msp1 clears mistargeted TA proteins from OMM by transferring them from mitochondria to the ER (Chen et al., 2014; Matsumoto et al., 2019; Okreglak and Walter, 2014). However, the surveillance mechanism which removes mistargeted TA/SA proteins from ER is still unknown.

Here we show that CATP-8, the worm homolog of ATP13A1 (mammalian) and Spf1 (yeast), functions as a surveillance factor to clear mitochondrial TA/SA proteins which ectopically localize to ER. CATP-8 belongs to the family of P-type ATPases, which is a large group of evolutionarily conserved ion and lipid transporters (Palmgren and Nissen, 2011). The P-type ATPases are divided into five groups based on sequence homology (Axelsen and Palmgren, 1998). The P1-P3 subgroup members transport different types of cations, while P4 ATPases transport/flip phospholipid molecules (Paulusma and Oude Elferink, 2005). The P5 subgroup, which is further divided into P5A and P5B classes, is the least characterized class. Most of the progresses of P5A ATPase are from the yeast homolog of Spf1. It is reported that Spf1 regulates manganese transport into the ER (Cohen et al., 2013), and is involved in ER function and calcium homeostasis (Cronin et al., 2002). Furthermore, Spf1 mutant shows reduced ER ergosterol content and mislocalization of mitochondrial TA/SA proteins to ER (Krumpe et al., 2012). While the sequence of Spf1 is homologous to *catp-8* in worm and ATP13A1 in mammal, the functional conservation of this class of ATPase has not been tested.

We show that CATP-8 localizes to ER but not to mitochondria. In *catp-8* mutants, multiple mitochondrial TA/SA proteins ectopically localize to ER membrane in both muscle cells and neurons. Transient expression of CATP-8 under the heatshock promoter in *catp-8* mutant is sufficient to remove the ectopically localized mitochondrial SA/TA proteins from ER, suggesting it functions as a surveillance mechanism to clear mistargeted mitochondrial proteins. In *catp-8* mutant, neuronal ER breaks in PVD dendrites due to ectopic action of mitochondrial fission protein DRP-1. CATP-8 is also required for PVD dendrite arbor morphogenesis by regulating the level of dendrite receptor DMA-1.

## RESULTS

### Mitochondrial signal/tail-anchored proteins are mislocalized to ER in *catp-8* mutants

Accurate targeting of membrane proteins to specific organelle is essential for its identity and physiological functions. ER is the most expansive organelle in the cell and many non-ER proteins, especially mitochondrial OMM proteins, are prone to be mislocalized to the surface of ER (Vitali et al., 2018). To ask if there are mechanisms that prevent mislocalization of mitochondrial proteins to ER, we performed an unbiased forward genetic screening using a mitochondrial SA protein TOMM-20 (1-54AA)::GFP in *C. elegans* sensory neuron PVD (Figure 1A). In wild type PVD, TOMM-20(1-54AA)::GFP was localized to discrete puncta along the highly branched dendritic processes and the axon (Figures 1A and 1B), a pattern that reflects the stringent targeting of TOMM-20(1-54AA)::GFP to mitochondrial structures. In the PVD soma, TOMM-20(1-54AA)::GFP showed a nuclear-excluded tubular and round pattern in the cytosol (Figure 1C), consistent with the tubular/round mitochondria in the PVD cell body from serial electron microscopy reconstruction (Data not shown). From this screen, we isolated three mutants that caused ectopic localization of TOMM-20 (Figure S1A). In these mutants, TOMM-20(1-54AA)::GFP or full length TOMM-20::GFP were localized to both puncta as well as long tubules along the primary dendrite and small numbers of secondary dendrite which closely resembles the distribution pattern of ER in PVD dendrites (Figures 1B and S1B). We have previously reported that ER tubules are found in the entire primary dendrite and a small percentage of secondary dendrites of PVD (Figures 1A and 1D) (Liu et al., 2019). Indeed, when visualized together with an ER marker (mCherry::SP12), TOMM-20 did not colocalize with ER in the wild type but was localized to ER in the mutant (Figures 1C and 1D). In the soma, TOMM-20 localized to mitochondrial tubules as wild type animals, but also showed a more diffusive pattern with concentration in perinuclear area in the mutants, which also colocalized with the ER marker (Figure 1C). Along the dendrite, the co-localization between TOMM-20 and the ER marker was unequivocal in the mutant (Figure 1D). TOMM-20 is a signal-anchored protein with its mitochondrial targeting sequence at its amino terminus. To test if the localization of other mitochondrial proteins is also affected by this mutation, we examined the localization of the TA protein MIRO-1 (ortholog of MIRO), whose transmembrane domain is at the carboxyl terminus of its coding sequence. MIRO-1 showed mitochondrial localization pattern in wild type PVD but ectopic localization to ER in the mutant (Figures 1E and 1F), suggesting a more general defect in mitochondrial proteins targeting in this mutant. Furthermore, we found a dramatic localization defect of MIRO-1 in the muscles (Figure 1H). Strikingly, unlike the ordered, round or linear array-like mitochondria localization in wild type muscles, MIRO-1 displayed an additional web-like pattern including a clear perinuclear ring which colocolized with an ER marker in the mutant, highly suggestive of MIRO-1’s mis-localization to ER (Figure 1H). Another mitochondrial TA protein FIS-1 (ortholog of FIS1) also showed ectopic ER localization in the mutant (Figure S1D). We next examined the localization of another mitochondrial protein, FZO-1 (ortholog of mitofusin) with two transmembrane domains and found that it was correctly localized to mitochondria in the mutant without mislocalization to the ER (Figure S1E). Together, these data argue strongly that the corresponding gene is required for the stringent localization of TA/SA proteins but not all mitochondrial proteins.

**Figure 1.**
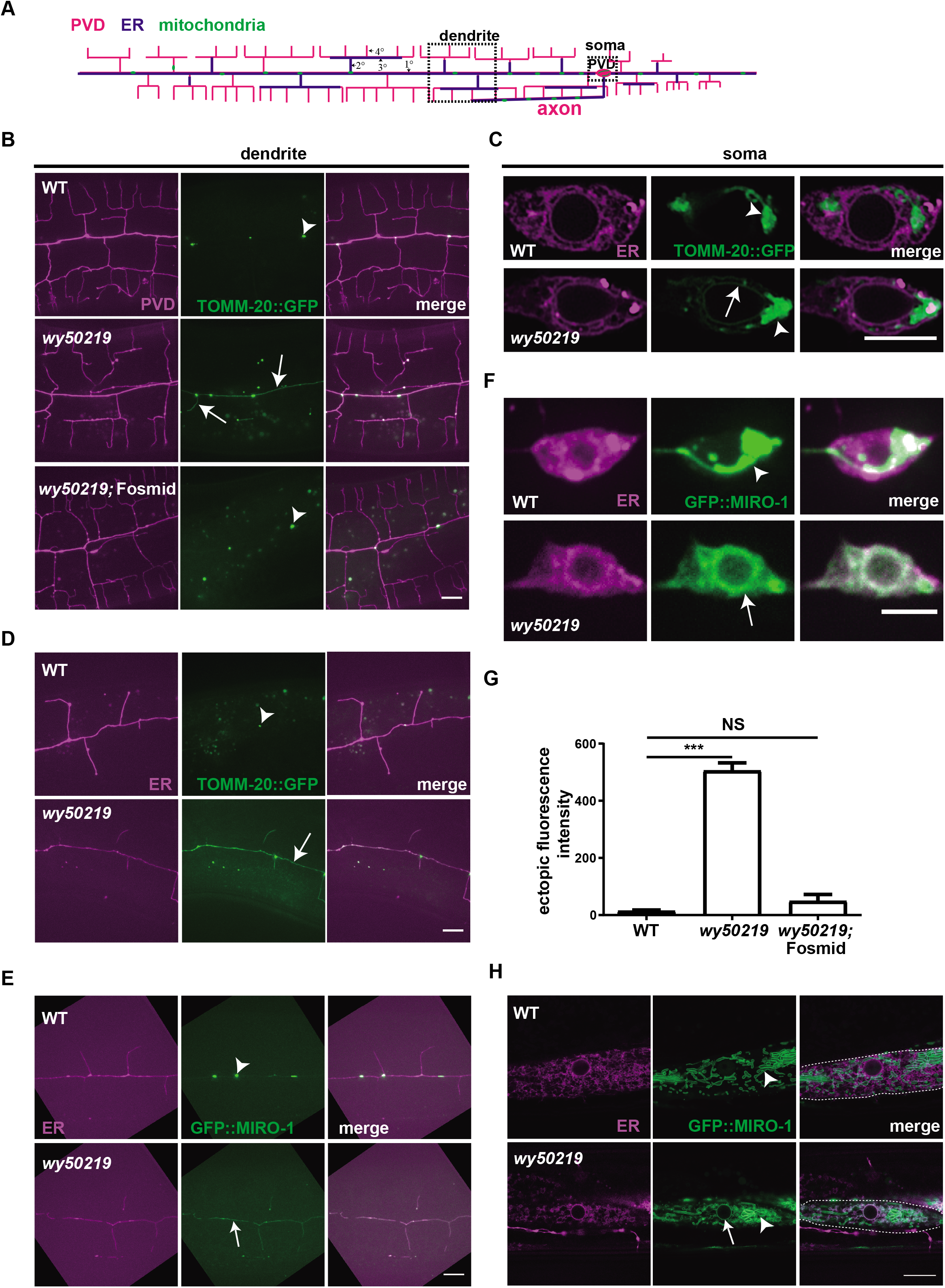
Mitochondrial signal/tail-anchored proteins are mislocalized to ER in *catp-8* mutants. **(A)** A schematic diagram showing PVD dendrites and mitochondrial distribution, ER pattern in the axon and dendrites of PVD. Magenta marks the PVD morphology with branched dendritic arbors containing ordered primary (1°), secondary (2°), tertiary (3°), and quaternary (4°) branches. Blue lines, which occupy part of the dendrites, indicate ER. Green dots along the neurites represent mitochondria. Anterior is to the left and dorsal is up. **(B)** Representative confocal images showing mitochondrial pattern in PVD dendrites in wild type, *wy50219*, and *wy50219;* Fosmid rescue strains. Magenta: PVD::mCherry. Green: PVD::TOMM-20(1-54AA)::GFP. Arrowheads indicate mitochondria and arrows show ectopic TOMM-20(1-54AA)::GFP. Scale bar: 10 μm. **(C-D)** Representative confocal images showing ER and mitochondrial pattern in PVD cell soma (C) and dendrites (D) in wild type and *wy50219* strains. Magenta marks ER. Arrowheads indicate mitochondria and arrows show ectopic TOMM-20(1-54AA)::GFP. Scale bar: 5 μm for (C) and 10 μm for (D). **(E-F)** Representative confocal images showing ER and GFP::MIRO-1 in PVD dendrite (E) and cell soma (F) in wild type and *wy50219*. Magenta marks ER. Arrowheads indicate mitochondria and arrows show ectopic GFP::MIRO-1. Scale bar: 10 μm for (E) and 5 μm for (F). **(G)** Quantification of ectopic TOMM-20(1-54AA)::GFP intensity in PVD dendrites in wild type, *wy50219*, and *wy50219;* Fosmid rescue strains. Data are shown as mean±SEM. One way ANOVA with Tukey correction. ***p<0.001. NS means not significant. n>=30 for each genotype. **(H)** Representative confocal images showing GFP::MIRO-1 in muscle of wild type and *wy50219*. One muscle cell is indicated by the dashed lines. Magenta marks ER. Arrowheads indicate mitochondria and arrows show ectopic GFP::MIRO-1. Scale bar: 10 μm.

In our genetic screen, all of the three isolated mutants fell into the same complementation group. Using single nucleotide polymorphorism (SNP) mapping and sequencing, we identified corresponding mutations in the *catp-8* gene. Transgenic expressing wild type copies of *catp-8* (Fosmid) in these mutants showed complete rescue of the TOMM-20 mislocalization phenotype (Figures 1B and 1G). These results suggest that CATP-8 prevents TA/SA mitochondrial proteins from localizing to ER and is hence required for stringent mitochondrial proteins targeting. Interestingly, a previous study showed that the yeast homolog of CATP-8, Spf1 is required for stringent targeting of TA/SA proteins to mitochondria, suggesting that this function is conserved in eukaryotes (Krumpe et al., 2012).

To test if endogenous level of mitochondrial protein is also affected by *catp-8* mutant, we constructed a single copy transgene TOMM-20::GFP in PVD by MOS1 technique (Frokjaer-Jensen et al., 2014). Despite of the low expression level, we observed clear GFP signal around the nuclei of PVD in *catp-8* mutant (Figure S1F), which is similar to an ER pattern (Liu et al., 2019).

### CATP-8 functions cell autonomously and localizes to ER

To test the tissue expression pattern of *catp-8*, we fused GFP with 2.5kb upstream sequence of the operon that contains *catp-8* and a second gene *mbf-1* (Figure 2A). The GFP signal was detected in PVD as well as several head and tail neurons, pharynx, body wall muscles and intestine, indicating that *catp-8* is broadly expressed in many tissues (Figure 2B).

**Figure 2.**
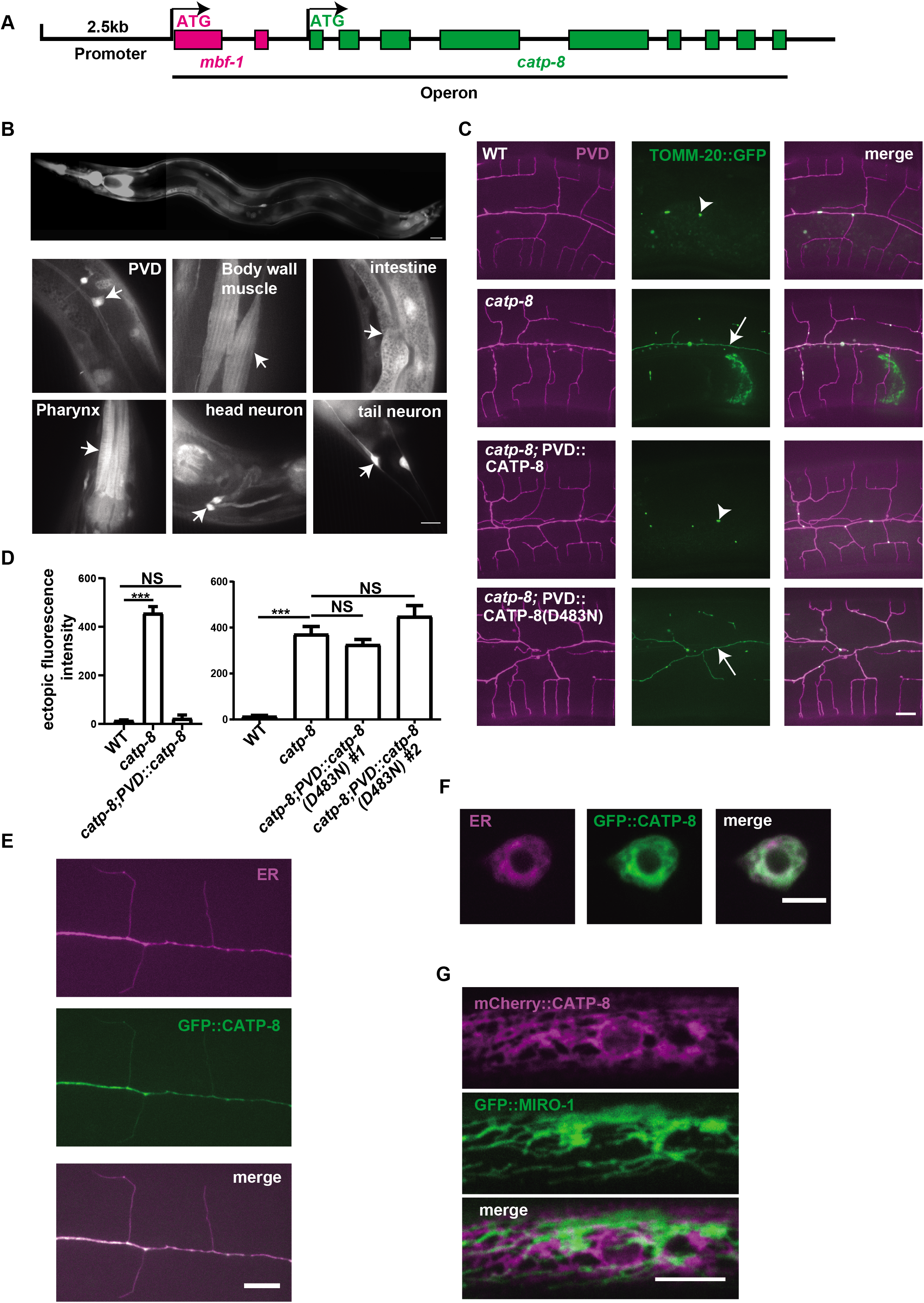
CATP-8 functions cell autonomously and localizes to ER. **(A)** A schematic diagram showing the operon, which includes *mbf-1* and *catp-8*. **(B)** The expression pattern of upstream 2.5kb sequence of *mbf-1* fused to GFP. Scale bar: 10 μm. **(C)** Representative confocal images showing mitochondrial pattern in PVD dendrites in wild type, *catp-8, catp-8; PVD::catp-8*, and *catp-8;* PVD::*catp-8*(D483N) rescue strains. Magenta: PVD::mCherry. Green: PVD::TOMM-20(1-54AA)::GFP. Arrowheads indicate mitochondria and arrows show ectopic TOMM-20(1-54AA)::GFP. Scale bar: 10 μm. **(D)** Quantification of ectopic TOMM-20(1-54AA)::GFP intensity in PVD dendrites in wild type, *catp-8, catp-8; PVD::catp-8*, and *catp-8;* PVD::*catp-8*(D483N) rescue (line1 and line2) strains. Data are shown as mean±SEM. One way ANOVA with Tukey correction. ***p<0.001. NS means not significant. n>=16 for each genotype. **(E)** Representative confocal images showing ER and GFP::CATP-8 pattern in PVD dendrite. Magenta marks ER. Scale bar: 10 μm. **(F)** Representative confocal images showing ER and GFP::CATP-8 pattern in PVD cell soma. Magenta marks ER. Scale bar: 5 μm. **(G)** Representative confocal images showing mCherry::CATP-8 and GFP::MIRO-1 pattern in muscle. Scale bar: 10 μm.

To test the cell autonomy of CATP-8, we expressed wild-type *catp-8* in PVD using the *ser-2P3* promoter, which drives expression in PVD and two head neurons. This wild type CATP-8 transgene fully rescued the TOMM-20 mislocalization phenotype in PVD (Figures 2C and 2D), indicating that CATP-8 functions cell autonomously. Next, we examined the subcellular localization of *catp-8* by creating a GFP fusion protein with CATP-8 and expressing this construct in PVD. GFP::CATP-8 showed striking colocalization with an ER marker, displaying a reticular pattern in the soma and a tubule-like pattern along the dendrite (Figures 2E and 2F). In the muscle cells, mCherry::CATP-8 also showed an ER pattern and did not show any colocalization with the mitochondrial marker GFP::MIRO-1 (Figure 2G). Together, these data suggest that CATP-8 functions cell autonomously and is localized to ER.

CATP-8 is a P5A type ATPase, and the ATPase activity of its yeast homolog Spf1 is essential for its function (Cohen et al., 2013). To test if the ATPase activity of CATP-8 is required for its function in PVD, we introduced a mutation (D483N) to disrupt the ATPase domain of CATP-8 (Cohen et al., 2013). Two independent transgenic lines expressing the D483N plasmid could not rescue the TOMM-20(1-54AA)::GFP mislocalization phenotype in *catp-8* mutant (Figures 2C and 2D), indicating the ATPase activity is essential for CATP-8 to safeguard mitochondrial proteins targeting.

### Ca^2+^ homeostasis are unlikely to explain the *catp-8* mutant phenotypes in PVD neuron

The yeast homolog of CATP-8, Spf1 has been implicated in regulating manganese transport into the ER and maintaining calcium homeostasis (Cronin et al., 2002). We speculated that the mitochondrial proteins targeting defects might be due to imbalance of cation in the ER lumen. To test this hypothesis, we measured ER calcium levels with a calcium sensor YC3.6 which was targeted to the ER lumen. YC3.6 is a genetically encoded fluorescent calcium sensor, which is a fusion protein that consists of a ECFP, calmodulin, the calmodulin-binding peptide M13 and cpVenus (YFP). (Nagai et al., 2004). We calculated the steady state fluorescence energy transfer (FRET) signal by quantifying the increase in CFP fluorescence after photobleaching YFP. With this method, we did not observe any obvious difference between wild type and *catp-8* mutant, suggesting that ER calcium is normal in *catp-8* mutant (Figures S2A and S2B). We also depleted or added 10x calcium in the diet to culture wild type and *catp-8* mutant. However, both treatments did not suppress or enhance the TOMM-20 mislocalization phenotype of *catp-8* mutant (Figure S2C). Together, these negative data do not support the notion that CATP-8 regulates mitochondrial proteins localization through controlling cation transport across the ER membrane. However, because of the limitation of our Ca^2+^ measurements, we cannot exclude the possibility that CAPT-8 plays a role in calcium homeostasis in ER.

### Rapid clearance of mistargeted TOMM-20 from the ER by heat shock expression of CATP-8

In wild type animals, TOMM-20 and other mitochondrial proteins displayed tight punctate staining pattern along the dendrite with undetectable fluorescence in the cytosol (Figures 1B, S1D, and S1E). This stringent mitochondria localization suggests that TOMM-20 is rapidly targeted to mitochondria after protein synthesis. To further understand the action of CATP-8, we considered two possibilities. The first idea is that CATP-8 prevents ectopic protein insertion into ER shortly after the protein is synthesized and before the proteins are fully inserted into the ER membrane. Alternatively, CATP-8 might remove the mis-targeted proteins from ER after they are fully inserted into the ER membrane.

To distinguish these two possibilities, we expressed CATP-8 under the control of the *hsp-16.48* (heat shock) promoter, in *catp-8* mutant. The *catp-8* mutant and the *catp-8;* P*hsp-16.48*:: CATP-8 animals were cultured at 15°C until young adults. These animals were then exposed to 30°C heatshock for one, two, or three hours. In the absence of heatshock, TOMM-20(1-54AA)::GFP fluorescence is detected on the ER in PVD cell body as evident from the nuclear envelope pattern (Figure 3A). Transgenic animals showed robust rescue of this phenotype after three hours of heatshock, an effect that is completely dependent on the transgene (Figures 3A and 3B). Similar rescue effects were observed in PVD dendrite, where the continuous ER tubule staining pattern within the primary and secondary dendrites became punctate mitochondria pattern after heatshock (Figures 3A and 3C). Interestingly, rescue of the TOMM-20 targeting phenotype can be detected even after just one hour of heatshock (Figures 3A and 3B). This rapid recovery hints that CATP-8 might be able to remove mistargeted TOMM-20 from ER.

**Figure 3.**
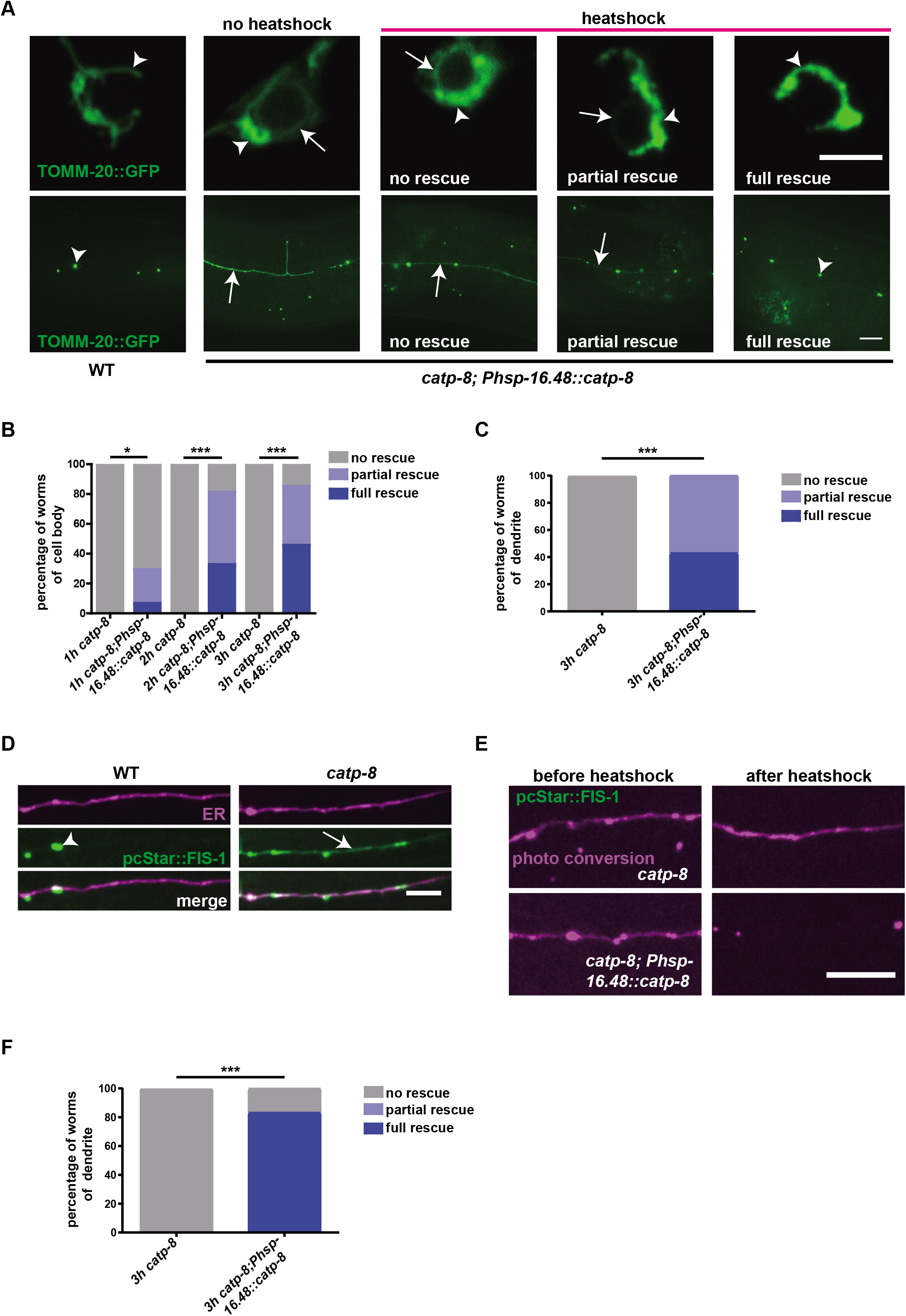
Efficient clearance of mistargeted TOMM-20 and FIS-1 from the ER by heat shock expression of CATP-8. **(A)** Representative confocal images showing mitochondrial pattern in PVD cell soma and dendrites in wild type and *catp-8; Phsp-16.48::catp-8* before and after heatshock. The heat shock rescue results are divided into three classes: no rescue, partial rescue, and full rescue, and the representative pictures are shown. Arrowheads indicate mitochondria and arrows show ectopic TOMM-20(1-54AA)::GFP. Scale bar: 5 μm for the first line and 10 μm for the second line. **(B)** Quantification of the percentage of ectopic TOMM-20(1-54AA)::GFP phenotype in PVD cell soma in *catp-8* and *catp-8; Phsp-16.48::catp-8* with heat shock (1h, 2h, and 3h). Fisher’s Exact Test, *p<0.05. ***p<0.001. n>=21 for each genotype. **(C)** Quantification of the percentage of ectopic TOMM-20(1-54AA)::GFP phenotype in PVD dendrites in *catp-8* and *catp-8; Phsp-16.48::catp-8* with heat shock for 3h. Fisher’s Exact Test, ***p<0.001. n>=25 for each genotype. **(D)** Representative confocal images showing ER and pcStar::FIS-1 in PVD dendrites in wild type and *catp-8* strains. Magenta marks ER. Arrowhead indicates mitochondria and arrow shows ectopic pcStar::FIS-1. Scale bar: 5 μm. **(E)** Representative confocal images showing pcStar::FIS-1 before and after heat shock for 3h in PVD dendrites in *catp-8* and *catp-8; Phsp-16.48::catp-8* worms. Scale bar: 5 μm. **(F)** Quantification of the percentage of ectopic pcStar::FIS-1 phenotype in PVD dendrites in *catp-8* and *catp-8; Phsp-16.48::catp-8* with heatshock for 3h. Fisher’s Exact Test, ***p<0.001. n>=18 for each genotype.

To test this directly, we repeated the above-mentioned experiment but replaced TOMM-20(1-54AA)::GFP with TOMM-20(1-54AA)::pcStar, which is a photoconvertible fluorescent protein (Zhang et al., 2019). Before heat shock, the animals were exposed to blue laser for 10 seconds to convert the fluorescence of pcStar from green to red (Figure S3A). And the red fluorescent signal indicates TOMM-20 proteins that are already mislocalized to ER at the time of photoconversion. We found that heat shock in the transgenic strain expressing P*hsp-16.48*::CATP-8 could decrease the red TOMM-20(1-54AA)::pcStar signal from perinuclear ER (Figures S3A and S3B). This result indicates that CATP-8 can remove mistargeted protein from ER to achieve stringent targeting of mitochondrial signal-anchored protein. To further test this hypothesis, we used pcStar::FIS-1 to repeat the heatshock experiment of PVD dendrite in the *catp-8* mutant animals. FIS-1 is a tail-anchored mitochondria outer membrane protein involved in mitochondria fission (Loson et al., 2013). Firstly, we co-expressed pcStar::FIS-1 and ER marker mCherry::SP12 and pcStar::FIS-1 colocalized well with ER marker, suggesting FIS-1 ectopically localized to ER in *catp-8* mutant (Figure 3D). Next, we repeat the heatshock experiments combined with photoconversion. We found that heatshock-expression of CATP-8 reduced or eliminated the ER localized pcStar::FIS-1 efficiently three hours after heatshock (Figures 3E and 3F), consistent with the TOMM-20::pcStar data.

### CATP-8 clears mistargeted TOMM-20 from the ER independent of the ERAD pathway

CATP-8 is P5-type ATPase with multiple transmembrane domains. Previous studies suggest that this family of ATPase functions as cation transporters. It is less likely that CATP-8 can directly bind to and extract ectopic proteins from ER. ER-associated degradation (ERAD) is a robust mechanism to degrade misfolded luminal or integral membrane proteins from ER. The essential players of this pathway includes Hrd1, Der1, Usa1, Doa10, and CDC48 complex, which function together to extract and degrade misfolded proteins (Brodsky, 2012).

We hypothesize that CATP-8 might be a regulator of ERAD. To test this hypothesis, we examined TOMM-20(1-54AA)::GFP marker in ERAD mutants including *sel-1, sel-11, hrdl-1, ubc-7, marc-6, cdc-48.1, cdc-48.2* (Miedel et al., 2012; Wu and Rapoport, 2018) and found that none of the single mutant showed the TOMM-20 mislocalization phenotype (Figure S4A). Then we constructed double mutants between *sel-11*, *marc-6*, *cdc-48.1* and *catp-8* and found that none of the ERAD mutations modified the *catp-8* single mutant phenotype (Figure S4B). The AAA-ATPase CDC-48 complex extracts ERAD substrates from ER (Ye et al., 2001). There are two orthologs in *C. elegans, cdc-48.1* and *cdc-48.2*, and double mutants of them were lethal. Then we overexpressed dominant negative constructs of *cdc-48.1* (E311Q&E584Q) or *cdc-48.2*(E310Q&E583Q) in the *cdc-48.2* or *cdc-48.1* mutant background, respectively. However, neither of them caused the mistargeting phenotype (Figure S4A). Next, we overexpressed wild-type CDC-48.1 alone or together with CDC-48.2 in *catp-8* mutant, but did not observe any suppression of the *catp-8* phenotype (Figure S4C). Therefore, we could not find any evidence that CATP-8 functions through the ERAD pathway to remove mitochondrial proteins from ER.

### Ectopic recruitment of DRP-1 to ER causes ER fragmentation in *catp-8* mutants

Mislocalization of multiple mitochondria proteins onto ER is suggestive erosion of ER organelle identity in the *catp-8* mutants. What are the consequences of mistargeting of the mitochondrial proteins? Like in mammalian neurons, the ER in PVD is a continuous membrane system spanning the nuclear envelope, a tubular network in the soma and tubules in the primary and some secondary/tertiary dendritic branches (Figure 4A) (Liu et al., 2019). In *catp-8* mutants, when we observed the mislocalized TOMM-20(1-54AA)::GFP or GFP::MIRO-1 on ER, we noticed that the ER staining along the neurite were often discontinuous, suggesting that the ER tubules might be broken in PVD dendrites (Figures 1D and 1E). To verify this phenotype, we crossed the *catp-8* mutant to an ER marker in PVD and it showed robust ER breakage phenotype (Figures 4A and 4B), indicating that CATP-8 is required to maintain the continuous ER morphology.

**Figure 4.**
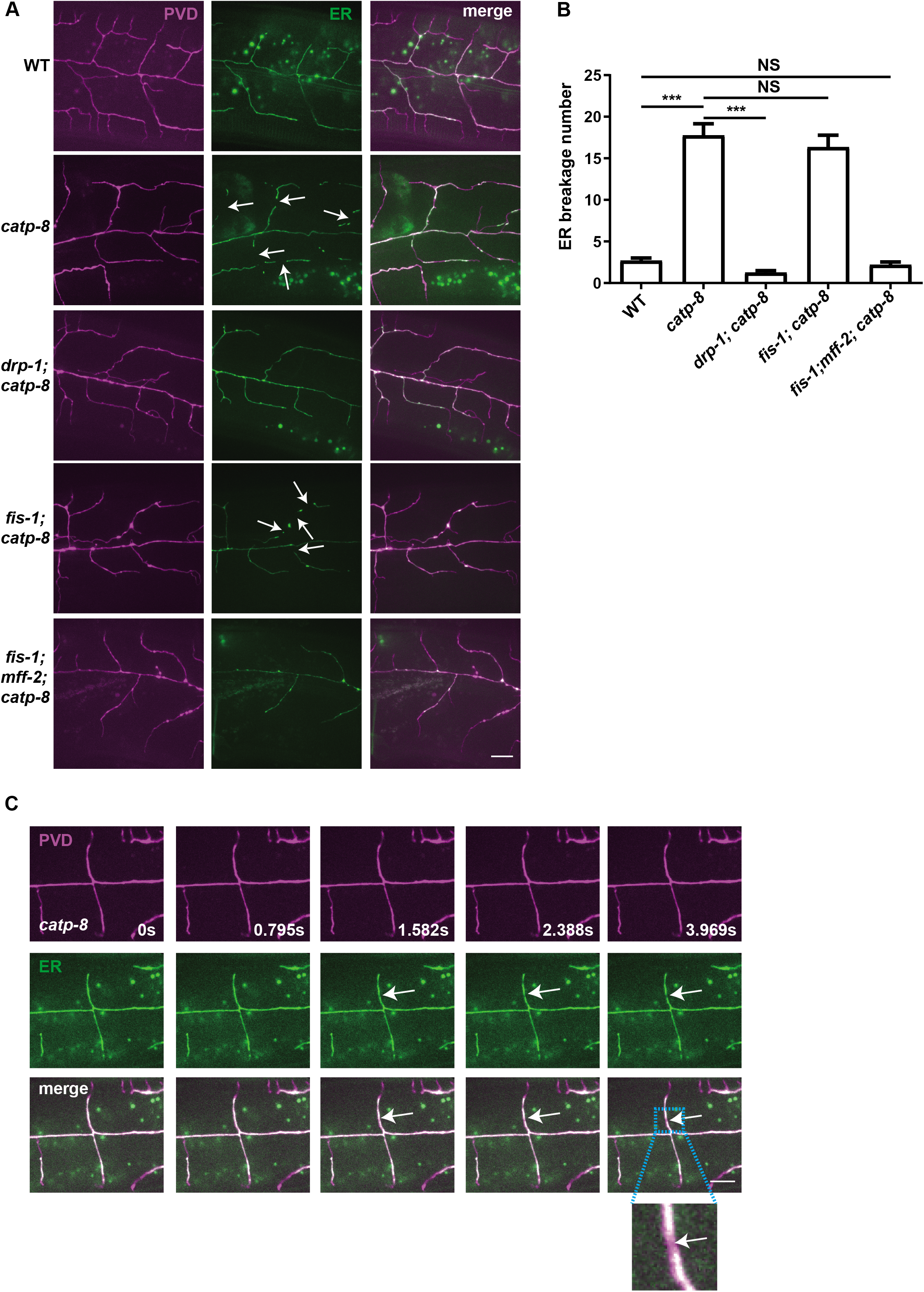
Ectopic recruitment of DRP-1 to ER causes ER breakage in *catp-8* mutants. **(A)** Representative confocal images showing ER pattern in PVD dendrites in wild type, *catp-8, drp-1; catp-8, fis-1; catp-8, and fis-1; mff-2; catp-8* mutants. Magenta: PVD::mCherry. Green shows ER. Arrows show breakage of ER. Scale bar: 10 μm. **(B)** Quantification of ER breakage phenotype in PVD dendrites in wild type, *catp-8, drp-1; catp-8, fis-1; catp-8, and fis-1; mff-2; catp-8* mutants. Data are shown as mean±SEM. One way ANOVA with Tukey correction. ***p<0.001. NS means not significant. n>=26 for each genotype. **(C)** Representative time-lapse images showing ER breakage event in the dendrite of PVD. Magenta marks the PVD dendrite and green shows ER. Arrows show the fragmentation site. Inset indicates the magnification of the fragmentation site. Scale bar: 10μm.

To understand how ER breaks, we performed time-lapse imaging on labeled ER in PVD dendrites in *catp-8* mutants. Certain sites of ER tubule became thinner through an apparent constriction process before breaking into two ends, which is a reminiscence of mitochondrial fission (Figure 4C). The dynamin related protein DRP-1 is recruited to mitochondrial surface to induce mitochondrial fission (Labrousse et al., 1999). So we hypothesize that ectopic DRP-1 localization to ER might be responsible for ER breakage. To test this hypothesis, we constructed the *drp-1; catp-8* double mutant. Remarkably, the double mutant showed complete suppression of the ER fragmentation phenotype of the *catp-8* single mutant (Figures 4A and 4B), indicating that DRP-1 is ectopically recruited to ER and causes ER fission in the absence of CATP-8. Since DRP-1 is a cytoplasmic protein, we next ask what is responsible for the ectopic action of DRP-1. For mitochondria fission, DRP1 is recruited to mitochondria by OMM protein Fis1 or Mitochondrial Fission Factor (Mff) (Loson et al., 2013). Indeed, we found that FIS-1 is ectopically localized to ER in the *catp-8* mutant (Figure S1D). However, a *fis-1* mutation did not suppress the ER fragmentation phenotype (Figures 4A and 4B), indicating that additional factors involved in recruiting DRP-1 to the ER to cause ER fragmentation. It is reported in yeast and the mammalian system that multiple proteins including *fis-2, mff-1*, and *mff-2* might serve as DRP-1 recruiters for mitochondria fission. To further test if DRP-1’s ectopic recruitment to ER is responsible for ER fragmentation in *catp-8* mutant, we generated *fis-1 mff-2 catp-8* triple mutants and found that the ER fragmentation phenotype of *catp-8* mutant was completely suppressed (figures 4A and 4B). These data suggest that *fis-1, mff-2* cooperatively recruit DRP-1 to ER in *catp-8* mutant. Together, these data argue strongly that mistargeting mitochondria proteins to ER leads to ectopic activation of DRP-1 and ER fragmentation.

### CATP-8 is essential for PVD dendrite morphogenesis through modulating DMA-1 level

*catp-8* mutants showed PVD dendrite defect, indicating that CATP-8 is required for PVD dendrite morphogenesis. In *catp-8*(*wy50219*) mutant, the total number of both secondary and quaternary branches were greatly reduced leading to a much reduced dendritic arbor (Figures 5B, 5D, and 5E). The phenotype was obvious at the 4^th^ larva stage (L4) and did not become stronger in Day3 (3 days after L4) adults (Figure S5), indicating the defect is developmental but not degenerative.

**Figure 5.**
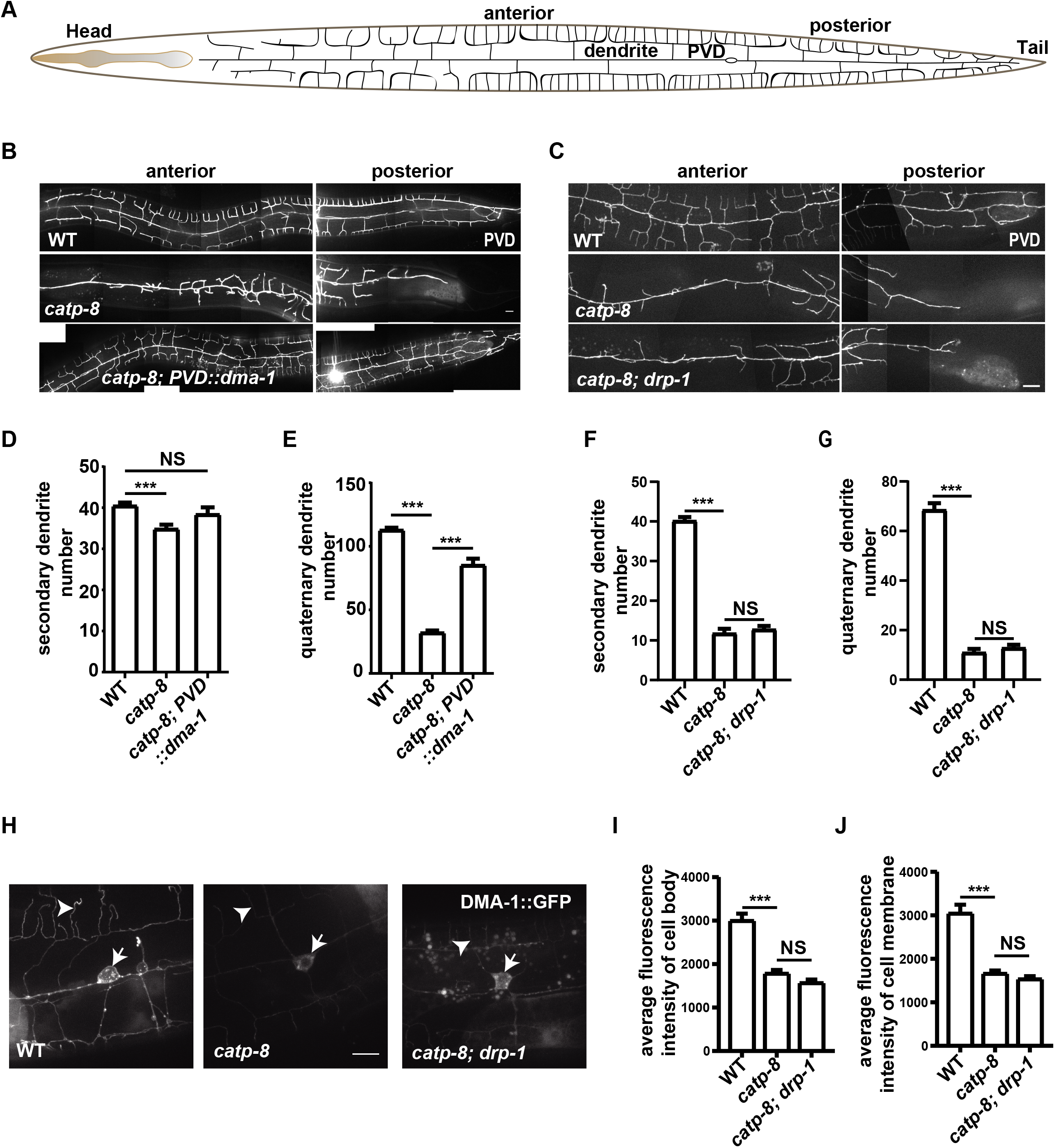
CATP-8 is essential for PVD dendrite morphogenesis through modulating DMA-1 level. **(A)** A schematic diagram showing PVD dendrites. **(B)** Representative images showing PVD anterior and posterior dendrite morphology in wild type, *catp-8, catp-8; PVD::dma-1* rescue strains. Scale bar: 10μm. **(C)** Representative images showing PVD anterior and posterior dendrite morphology in wild type, *catp-8, catp-8; drp-1* strains. Scale bar: 10μm. **(D-E)** Quantification of PVD secondary (D) and quaternary (E) dendrites phenotype in wild type, *catp-8, catp-8; PVD::dma-1* rescue strains. Data are shown as mean±SEM. One way ANOVA with Tukey correction. ***p<0.001. NS means not significant. n>=37 for each genotype. **(F-G)** Quantification of PVD secondary (F) and quaternary (G) dendrites phenotype in wild type, *catp-8, catp-8; drp-1* strains. Data are shown as mean±SEM. One way ANOVA with Tukey correction. ***p<0.001. NS means not significant. n>=32 for each genotype. **(H)** Representative confocal images showing DMA-1::GFP in PVD cell soma and dendrites in wild type, *catp-8, catp-8; drp-1* strains. Arrows show DMA-1::GFP in cell membrane and arrowheads indicate DMA-1::GFP in PVD dendrites. Scale bar: 10μm. **(I-J)** Quantification of DMA-1::GFP intensity of cell body (I) and cell membrane (J) in wild type, *catp-8, catp-8; drp-1* strains. Data are shown as mean±SEM. One way ANOVA with Tukey correction. ***p<0.001. NS means not significant. n>=21 for each genotype.

DMA-1, a single transmembrane leucine-rich repeat protein, is the adhesion receptor in PVD to build the PVD specific dendritic arbor (Dong et al., 2013; Liu and Shen, 2011; Salzberg et al., 2013; Zou et al., 2016). We ask if the dendrite morphology defect in *catp-8* mutant is due to DMA-1 level or localization change. We constructed a translational fusion DMA-1::GFP (GFP inserted 3 amino acids after the transmembrane domain) strain and compared the fluorescence of wild type to that of the *catp-8* mutant. The GFP signal was greatly reduced both along the entire PVD dendritic arbor and in the cell body (Figures 5H-J). Surprisingly, overexpression of DMA-1 rescued the PVD dendrite phenotype of *catp-8* mutant (Figures 5B, D, and E), indicating CATP-8 regulates PVD dendrite morphogenesis primarily through modulating DMA-1 level. To test if *catp-8* is required for general secretory/sorting proteins, we examined two other PVD membrane/secreted proteins in *catp-8* mutant background. HPO-30 is a four-pass transmembrane protein which functions in PVD as a co-receptor of DMA-1 to establish dendritic arbors (Zou et al., 2018) and NLP-21 is a neuropeptide (Konop et al., 2015). We found no difference between wild type and the *catp-8* mutant (Figures S6A and S6B), indicating that *catp-8* is not required for biogenesis of general transmembrane or secreted protein. Instead, it is required for specific proteins like DMA-1.

To test whether the ER fragmentation is the reason for reduced DMA-1 level in the *catp-8* mutant, we examined if the *drp-1* mutation could suppress the DMA-1 reduction in the *catp-8* mutant because the *drp-1* mutation indeed suppressed the ER breakage defects. *catp-8; drp-1* double mutants showed similarly severe PVD dendrite defects as the *catp-8* single mutant (Figures 5C, F, and G). Similarly, the *drp-1* mutation also did not change the DMA-1 expression level in the *catp-8* mutant (Figures 5H-J), indicating dendrite defect is not caused just by ER fragmentation.

In summary, CATP-8 functions in ER to clear ectopic mitochondrial TA/SA proteins. Mistargeting of mitochondrial proteins to ER causes defects in ER integrity and maybe reduced level of specific protein which causes dendrite branching defect.

## DISCUSSION

Signal-anchored (SA) proteins and tail-anchored (TA) proteins are targeted to ER and mitochondria where they play important functions for these organelles. The insertion of ER-destined TA proteins requires the GET pathway and the EMC complex. While the molecular mechanism to insert mitochondria-destined TA proteins have not been identified. ER-destined TA proteins mislocalize to mitochondria when the GET pathway is defective, arguing that the GET pathway also provides the specificity of TA protein targeting. It has also been described that the mitochondrial membrane protein Msp1, an AAA-ATPase, functions as surveillance mechanism to remove and initiate degradation of mistargeted tail-anchored proteins on OMM. However, similar surveillance mechanism on ER has not been found. Such ER surveillance might be important because data support the notion that the efficient GET pathway can increase the risk of OMM proteins to be mistargeted to the ER (Vitali et al., 2018).

Here, we report that a P5A type ATPase CATP-8 is a component of the ER surveillance pathway to clear mistargeted mitochondrial TA/SA proteins. First, CATP-8 localizes to ER specifically. Second, mitochondrial TA/SA proteins accumulate on ER surface in *catp-8* mutant. Third, transient expression of wild type CATP-8 protein to *catp-8* mutant rapidly clears existed mistargeted mitochondrial proteins from ER.

The membrane-anchored AAA-ATPase Msp1 binds directly with ectopic substrates and extracts them from OMM. CATP-8 is not an AAA-ATPase and has small chance to be able to extract the mistargeted proteins directly. However, a couple of recent studies did indicate that non-AAA ATPases can disassembly protein complexes (Demoinet et al., 2007; Gouridis et al., 2019). The most likely candidate is ERAD, which extracts and degrades unfolded ER proteins. However, blocking ERAD with various genetic approaches in worm did not show any mitochondrial SA protein targeting phenotypes (Figure S4A), suggesting that CATP-8 clears ectopic OMM SA proteins through an ERAD independent manner. It is plausible that another AAA-ATPase on ER might function together with CATP-8 to extract ectopic OMM SA proteins from ER.

Studies of the yeast homolog Spf1 of CATP-8 have shown that Spf1 is required for proper targeting of mitochondria TA/SA proteins (Krumpe et al., 2012). It was proposed that Spf1 controls the ergosterol level on ER membrane, which then affects the targeting specificity of TA/SA proteins. This model suggests that CATP-8 indirectly affects the lipid composition of ER membrane, which would impact the insertion of TA/SA proteins. This model predicts that CATP-8 should only be able to prevent newly synthesized mitochondrial proteins to be inserted onto ER but should not be able to remove the existed ectopic mistargeted proteins. The data from *C. elegans* PVD neuron argues for a different model. The *C. elegans* genome does not contain many key enzymes for ergosterol synthesis (Vinci et al., 2008), suggesting that CATP-8 might not function through regulating the key enzyme for sterol synthesis. In addition, transient expression of CATP-8 was sufficient to rapidly remove the ectopic proteins in PVD neurons (Figures 3A-F, Figures S3A-B). The rescuing effect could be observed for as short as one hour after heatshock. Considering the time for protein synthesis, it is very likely that this resucing effect is due to removal of ectopic proteins by a CATP-8-dependent mechanism which does not just work on newly synthesized proteins.

What is the physiological consequence of mistargeting mitochondria proteins to ER? We observed at least two different phenotypes in the *catp-8* mutant. First, the ER tubules in the PVD dendrite showed frequent breakage in the *catp-8* mutant. Unlike ER which is a continuous membrane network, mitochondria undergo fission and fusion. Mechanistically, mitochondria fission is mediated by dynamin like protein DRP-1, which is recruited to mitochondria by OMM protein Fis1 or Mitochondrial Fission Factor (Mff) (Loson et al., 2013). Indeed, we found that FIS-1 was ectopically localized to ER in the *catp-8* mutant. Furthermore, a mutation in *drp-1* completely suppressed the ER breakage phenotype in the *catp-8* mutant. However, a mutation in *fis-1* did not suppress the ER breakage phenotype, suggesting redundant factors recruit DRP-1 to ER. By constructing *catp-8 fis-1 mff-2* triple mutants, we found that the ER fragmentation phenotype of *catp-8* was completely suppressed in the triple mutants, arguing that *fis-1* and *mff-2* cooperatively recruit DRP-1 to ER in *catp-8* mutant. Together, these data suggest that ectopic targeting of FIS-1 and MFF-2 causes DRP-1 to be recruited to the ER surface, leading to ER breakage.

Second, we found that the *catp-8* mutants showed an oversimplified dendritic arbor, a phenotype caused by reduced growth of PVD dendrite during development. Our previous work shows that the neuronal adhesion receptor DMA-1 detects extracellular ligands to drive the dendrite growth and branching. Here, we found that DMA-1 level was dramatically reduced in the *catp-8* mutants. In addition, overexpression of DMA-1 in the *catp-8* mutants largely rescued the dendrite development, indicating that the dendrite defect is mainly caused by reduced DMA-1 level (Figures 5B, D, and E). The development and function of PVD also requires cell autonomous functions of other proteins including HPO-30 (Zou et al., 2018), the level of which was not affected by the *catp-8* mutation, suggesting that CATP-8 is required for the level of specific proteins. A companion study indeed shows that the signal sequence of *dma-1* causes *dma-1* to be expressed at a relatively low level and is responsible for its susceptibility to the *catp-8* mutation (Y. Z., personal communication). While the *drp-1* mutation suppressed the ER breakage phenotype of *catp-8* mutants, it did not suppress the PVD dendrite branching phenotype, indicating that the reduced DMA-1 level is not solely caused by ER breakage. Together, these data indicate that the ectopic localization of multiple mitochondrial proteins to ER impairs the morphology and likely impact physiological functions of ER, causing ER to break and might reduce the level of selective transmembrane proteins.

In summary, we report a surveillance factor CATP-8 on ER membrane to clear ectopic mitochondrial TA/SA proteins to maintain the ER identity and normal function. This mechanism is essential for neuronal ER morphology, transmembrane protein level and dendrite arbor formation.

## STAR □ METHODS

### Strains and genetics

N2 Bristol was used as the wild type strain. Animals were grown at 20°C under standard conditions on NGM plates using *Escherichia coli* strain OP50. Unbiased forward genetic screening were carried out using wild type animals, integrated with *ser-2*P3::TOMM-20(1-54AA)::GFP; *ser-2*P3::myri-mcherry transgenes to visualize PVD mitochondrial pattern, by using 50 mM ethyl methane sulfonate (EMS). Three mutants including *wy50219, wy50230*, and *wy50648* were isolated, and non-complementary test showed that they were different alleles of the same gene. SNP mapping were performed by standard protocol (Davis et al., 2005). All the strains used are listed in Supplementary Table 1.

CRISPR/Cas9-assisted knockout worms were generated following the described procedure (Dickinson et al., 2013). The guide RNA target sequence was selected according to the design tool (http://crispor.tefor.net/). Single-copy transgenes were generated using the miniMos-mediated single-copy insertion method (Frokjaer-Jensen et al., 2014). *gfp* was fused to *tomm-20* C-terminus, and then the *tomm-20::gfp* coding sequence were inserted into pWZ347 (a modified version of pCFJ909) with the *ser-2P3* promoter (~1.6 kb) and the *unc-54* 3’UTR, respectively. The single copy insertion plasmid pQQ50 (27 ng/ul), transposase plasmids pCFJ601 (50 ng/ul), and selective marker *Pord-1::rfp* (60 ng/ul) were co-injected into *unc-119(ed4)* worms. Integration lines were obtained by picking out the non-uncoordinated worms, and further confirmed by PCR and Sanger sequencing. The line *wySi50064*[*ser-2*P3::*tomm-20::gfp*] was selected for further characterization.

### Molecular cloning

Plasmids were constructed using the pPD95.77 backbone. PCR products were produced by Phusion DNA polymerase (New England Biolabs) or Q5 High-Fidelity DNA polymerase (New England Biolabs). In-Fusion PCR Cloning System was used to construct all the plasmids. All the plasmids used are listed in Supplementary Table 2.

### CaCl_2_ treatment

CaCl_2_ was dissolved in ddH_2_O to produce a 1M stock solution and were added into NGM media to achieve the final concentration of 1mM and 10mM. L4 stage animals were transferred to plates with different concentrations of CaCl_2_ to grow for 3 days. Then L4 stage offsprings were transferred to another new plate for 4 days, and the next generation animals were quantified at Day1 (one day after L4 stage) stage.

### Calcium Sensor and Acceptor-photobleaching-based FRET measurements

YC3.6 was used as ER calcium sensor which was fused with STC-1(1-29aa) at N-terminal and KDEL at C-terminal to target it to ER lumen. FRET measurements were done using laser scanning confocal imaging system (Olympus FV1000). FRET measurements were performed as previously described with some modifications (Fang et al., 2015). CFP was excited with 405nm laser and detected at 435nm-465nm. 515nm laser was used for YFP excitation and detected at 565nm-595nm. Acceptor photobleaching was achieved by scanning a region of interest (ROI) 50 times (scans at 2 μs pixel time) by using the 515 nm laser line at 100% intensity. Five donor and acceptor fluorescence images were taken before and after the acceptor photobleaching procedure to assess changes in donor and acceptor fluorescence. The FRET signal (efficiency) was calculated according to *E*FRET = (*I*_DA_-*I*_DB_)/*I*_DA_. I_DB_ was donor intensity of kinetochores before acceptor photobleaching. I_DA_ was donor intensity of kinetochores after acceptor photobleaching.

### Heatshock experiments

*catp-8* DNA was fused with heatshock promoter P*hsp-16.48* and injected into mutant animals in *ser-2*P3::TOMM-20(1-54AA)::GFP; *ser-2*P3::myri-mcherry transgenes.

Worms used for experiments were grown at 15°C to Day1 (one day after L4 stage).

Because of some leaky expression at 15°C, the TOMM-20 mistargeting phenotype in some mutants with P*hsp-16.48::catp-8* transgene were rescued even before heatshock and these animals were not used for this experiment. Animals with obvious TOMM-20 mistargeting phenotype were picked out and transferred to 30°C to induce heatshock expression of CATP-8 for 1h, 2h, and 3h, then worm numbers of full rescue, partial rescue, and no rescue were counted, respectively. Fluorescence conversion heatshock experiment was carried out by replacing GFP with pcStar (Zhang et al., 2019). Animals were put in M9 and mounted on 5%(w/v) agar pads. Those with obvious TOMM-20 mistargeting phenotype were photoactivated using 405nm laser for 10s under the 63X oil lense and then rescued to 30°C for 3h. Then the worms were quantified as mentioned above. pcStar::FIS-1 fluorescence conversion heatshock experiment was done using the same procedure as TOMM-20 but photoactivating using 405nm laser under 100X oil lense. Those worms, dead or florescence bleached from observation step, were not quantified.

### Fluorescent imaging, time-lapse imaging

*C. elegans* were anesthetized using 1 mg/ml levamisole in M9 buffer, and then mounted on 3% (w/v) agar pads. All images were taken by the spinning disk confocal imaging system (Yokogawa CSU-X1 Spinning Disk Unit) at the same condition except for special notification. Fig. 1C, 1H, and S1F images were taken by the Zeiss LSM980 with Airyscan 2 super-resolution imaging system. Fig. 2B were taken by the Zeiss imager M2. All images for figures and quantification were taken when worms are at Day1 stage except for special notification.

Dynamic images were recorded continually with 800ms interval for 3mins by the spinning disk confocal imaging system (Yokogawa CSU-X1 Spinning Disk Unit) at the same condition.

### Statistical analysis

To quantify TOMM-20 mistargeting phenotype, a 50μm long region near the vulva region was selected. Spinning disk confocal imaging system was used to take z-stack photos. Image J was used for maximum intensity stack projection to show ectopic flourescence at the dendritic branches including primary, secondary, tertiary, and quaternary dendrites at that region. Average florescence intensity (excluding big puntas that represent mitochondrial florescence) along PVD primary dendrite minus the average background florescence intensity of the same length next to the primary dendrite was calculated. To quantify ER breakage phenotype, the total number of break site at primary, secondary, and tetiary dendrites was calculated. For PVD dendritic phenotype, the total mumber of secondary and quternary dendrites was calculated.

For Fig. 3B, 3C, 3F, and S3B, worms that showed obvious ectopic ER localization in cell soma or dendrites were counted as no rescue; worms that showed clear mitochondrial pattern but still remained very dim ectopic ER localization were counted as partial rescue; worms that showed clear mitochondrial pattern with no ectopic ER localization were counted as full rescue. For Fig. 3C, all the PVD dendrites were quantified. For Fig. 3F, only the region of photoconvertion were quantified.

All the statistical analyses were performed using GraphPad Prism by one-way ANOVA with Tukey correction (three or more groups) or Student’s t test (two groups). Fisher’s Exact Test was used for Fig. 3 and Fig. S3. p>0.05, not significant, *p<0.05, **p< 0.01, and ***p<0.001. All the data were shown as mean±SEM. ImageJ and illustrator were used to process pictures.

## Supporting information

Supplementary tabel 1

Supplementary tabel 2

## ACKNOWLEDGMENTS

We are grateful to the *Caenorhabditis* Genetics Center for strains and Dr. M. Zhang and Dr. P. Xu for pcStar. We thank Yan Teng from Center for Biological Imaging (CBI), Institute of Biophysics, for technical support with confocal imaging and image analysis. This work was supported by grants from the National Key R&D Program of China (2017YFA0102601), the Strategic Priority Research Program of CAS (XDB37020302) and the National Natural Science Foundation of China (31771138) to X. Wang; as well as a grant from the Beijing Municipal Science & Technology Commission (Z181100001518001) and the National Natural Science Foundation of China (31829001) to K. Shen. K. Shen is an investigator of the Howard Hughes Medical Institute.

## AUTHOR CONTRIBUTIONS

K. S., X.M. W., and W. Z. initiated the project and guide the whole experiments. Q.Q and T. Z. did the experiments. Q.Q, X.M. W., and K. S. analyzed the data. X.M. W. and K. S. interpreted the data and wrote the manuscript. All authors reviewed the manuscript.

---

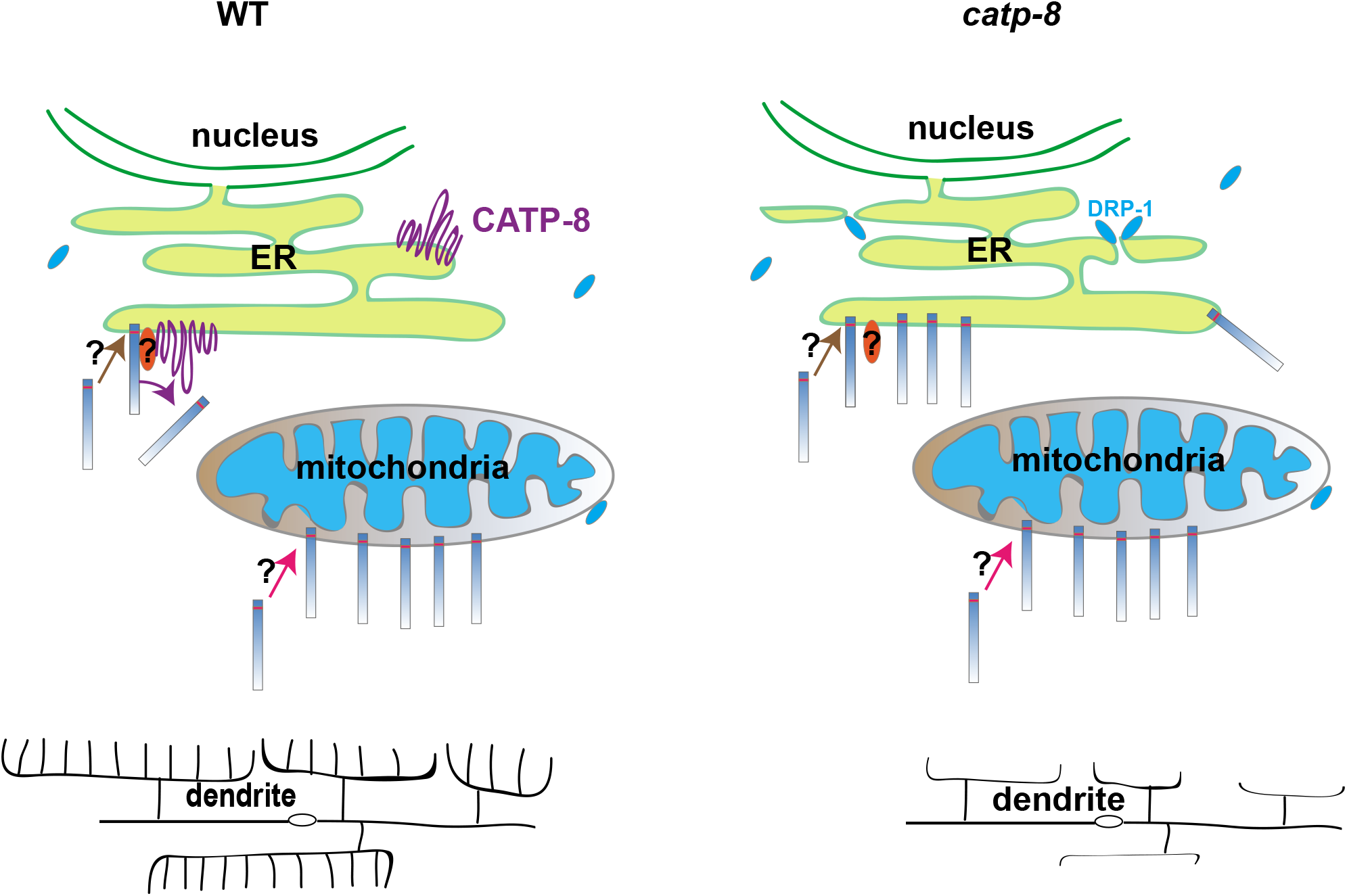

**Figure S1.**
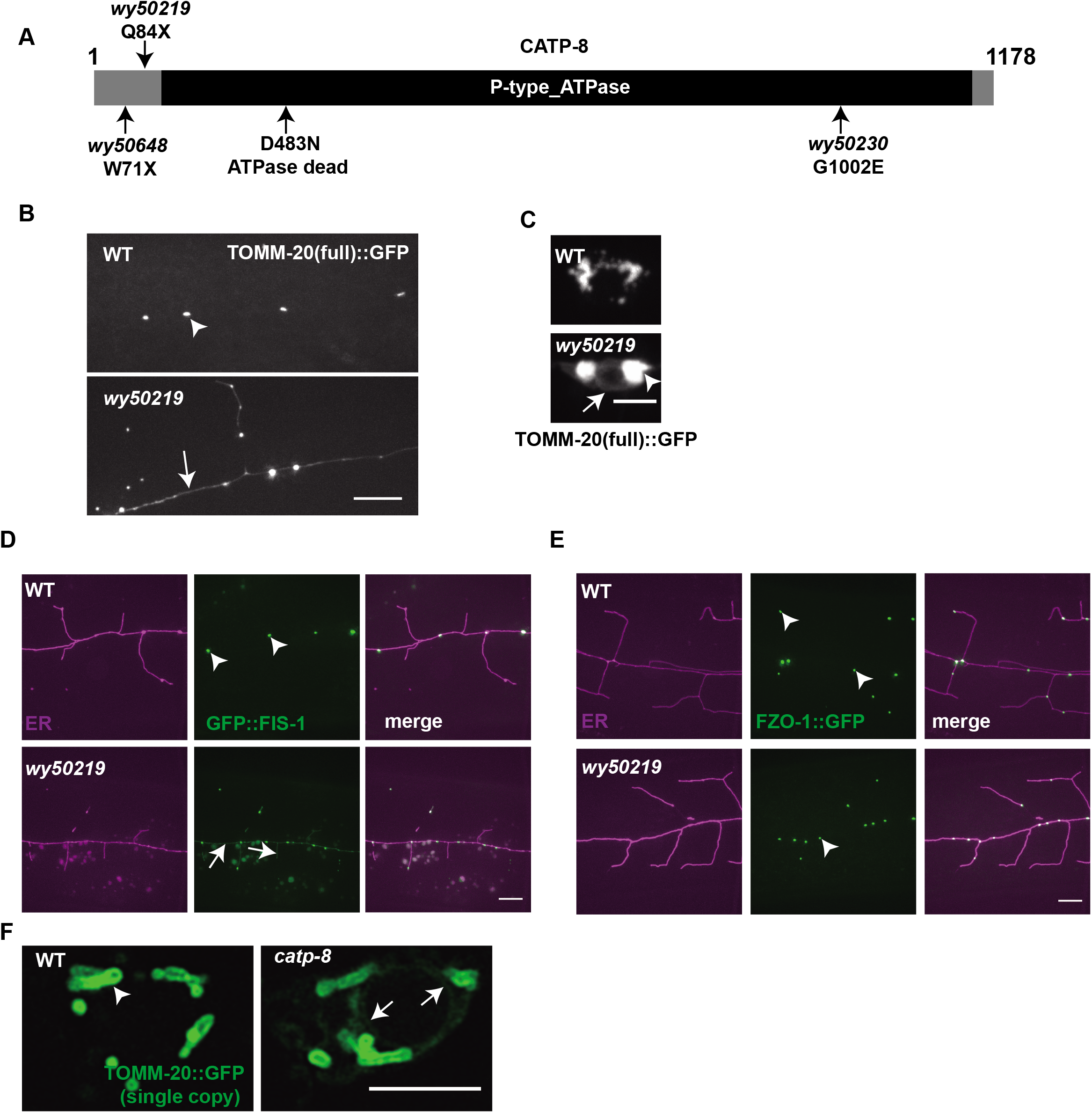
Mitochondrial signal/tail-anchored proteins are mislocalized to ER in *catp-8* mutants. **(A)** A schematic diagram showing alleles of *catp-8* isolated in our screen and the ATPase dead mutation site. **(B-C)** Representative confocal images showing mitochondrial pattern in PVD dendrites (B) and cell soma (C) in wild type and *catp-8* mutant. Arrowheads indicate mitochondria and arrows show ectopic TOMM-20(full)::GFP. Scale bar: 10 μm for (B) and 5 μm for (C). **(D)** Representative confocal images showing ER and mitochondrial pattern of GFP::FIS-1 in PVD dendrites in wild type and *catp-8* mutant. Arrowheads indicate mitochondria and arrows show ectopic GFP::FIS-1. Scale bar: 10 μm. **(E)** Representative confocal images showing ER and mitochondrial pattern using FZO-1::GFP in PVD dendrites in wild type and *catp-8* mutant. Arrowheads indicate mitochondria. Scale bar: 10 μm. **(F)** Representative confocal images showing mitochondrial pattern using single copy transgene of TOMM-20::GFP in PVD cell body in wild type and *catp-8* mutant. Arrowheads indicate mitochondria and arrows show ectopic TOMM-20::GFP. Scale bar: 5 μm.

**Figure S2.**
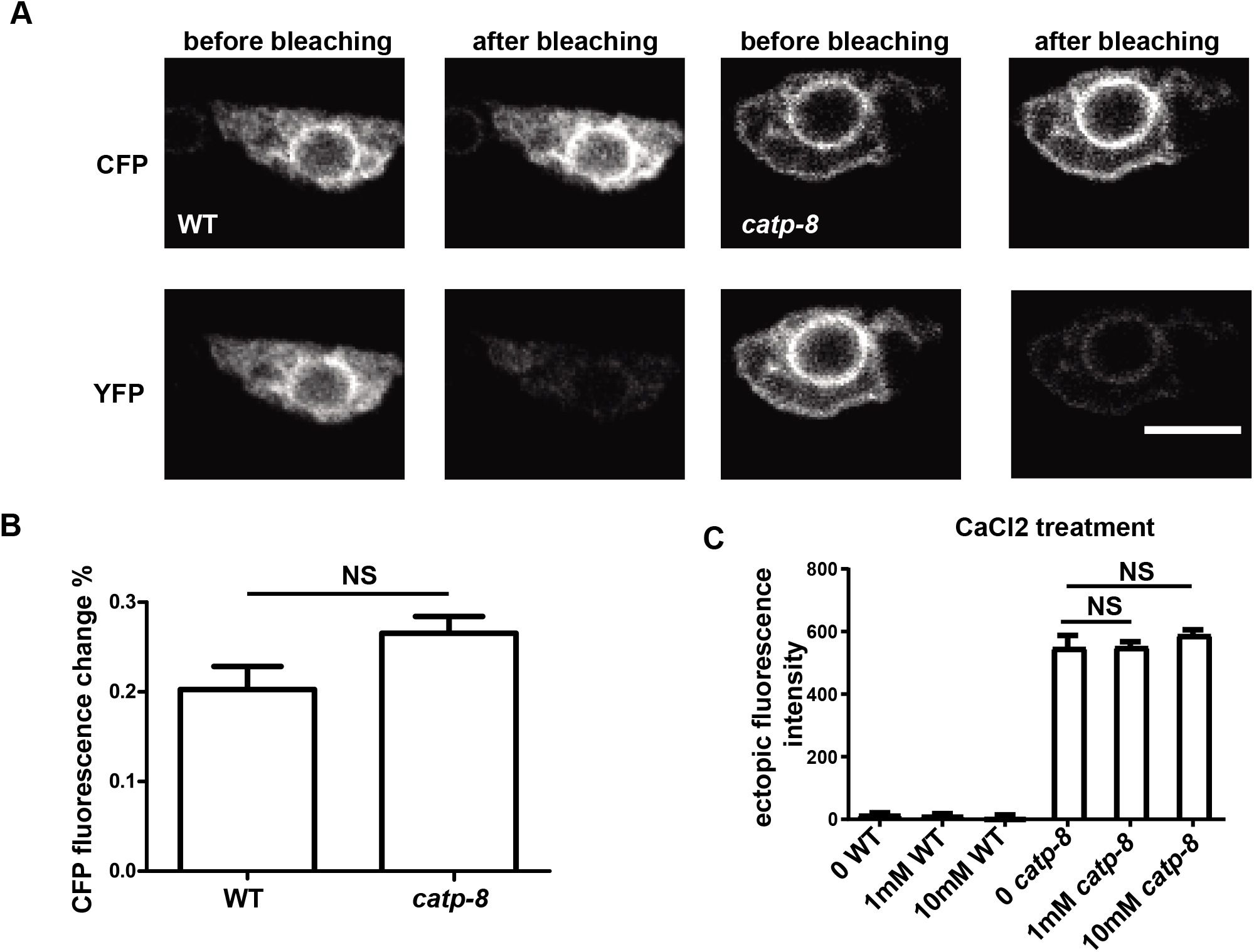
The cation transporting activity might not be essential for CATP-8 function. **(A)** Representative confocal images showing calcium sensor in PVD ER before and after bleaching YFP in WT and *catp-8* mutant. Scale bar: 5 μm. **(B)** Quantification of CFP fluorescence change ratio before and after bleaching YFP. Student’s t test, NS: not significant. n>=25 for each genotype. **(C)** Quantification of ectopic TOMM-20(1-54AA)::GFP intensity in PVD dendrites in wild type and *catp-8* mutant treated with different concentration of CaCl2. Data are shown as mean±SEM. One way ANOVA with Tukey correction. NS means not significant. n>=19 for each genotype.

**Figure S3.**
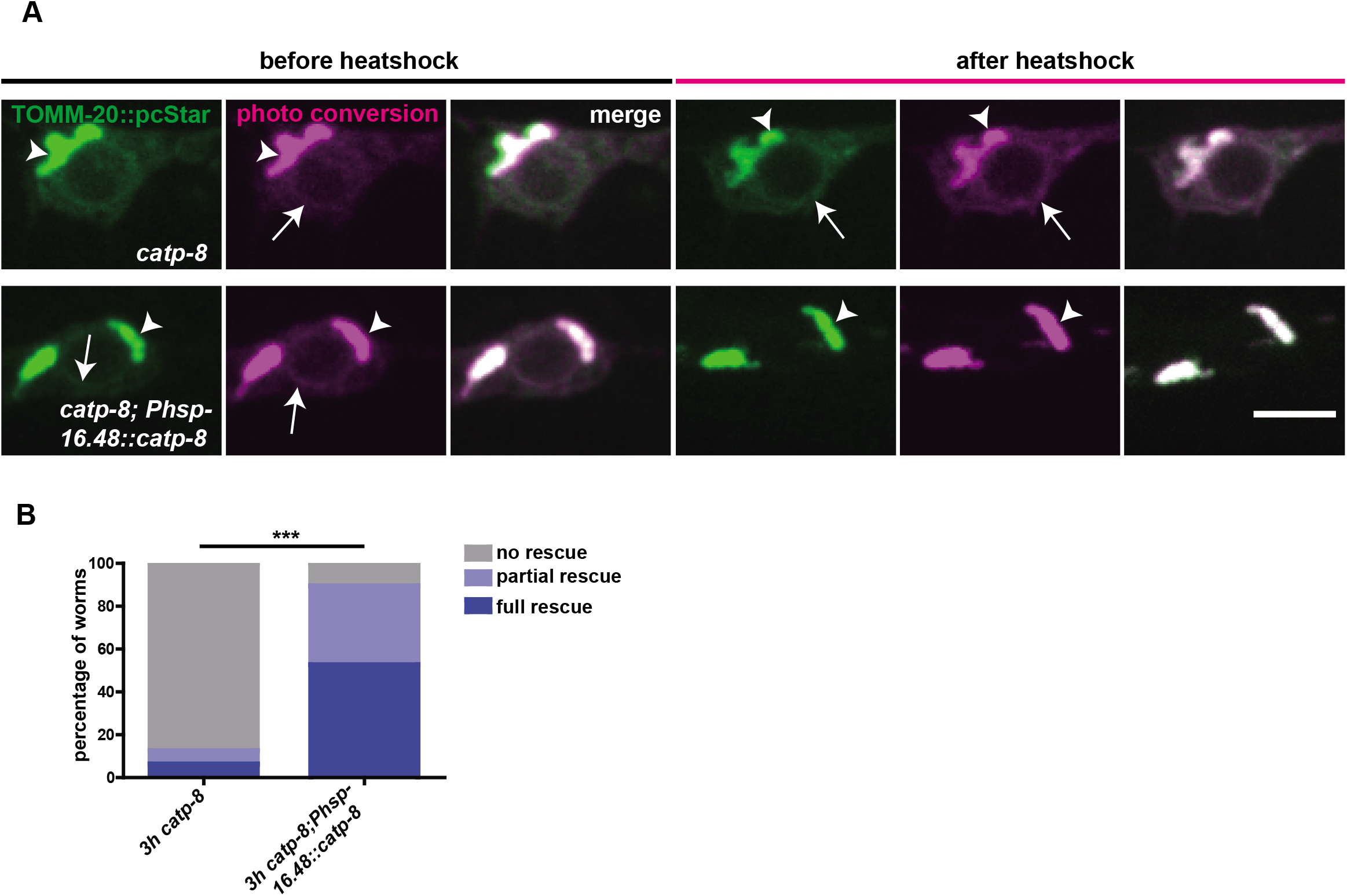
Efficient clearance of mistargeted TOMM-20 from the ER by heat shock expression of CATP-8 in cell soma. **(A)** Representative confocal images showing TOMM-20(1-54AA)::pcStar in PVD cell soma in *catp-8* and *catp-8; Phsp-16.48::catp-8* before and after heat shock for 3h. **(B)** Quantification of the percentage of ectopic TOMM-20(1-54AA)::pcStar phenotype in PVD cell soma in *catp-8* and *catp-8; Phsp-16.48::catp-8* after heatshock for 3h. Fisher’s Exact Test, ***p<0.001. n>=16 for each genotype.

**Figure S4.**
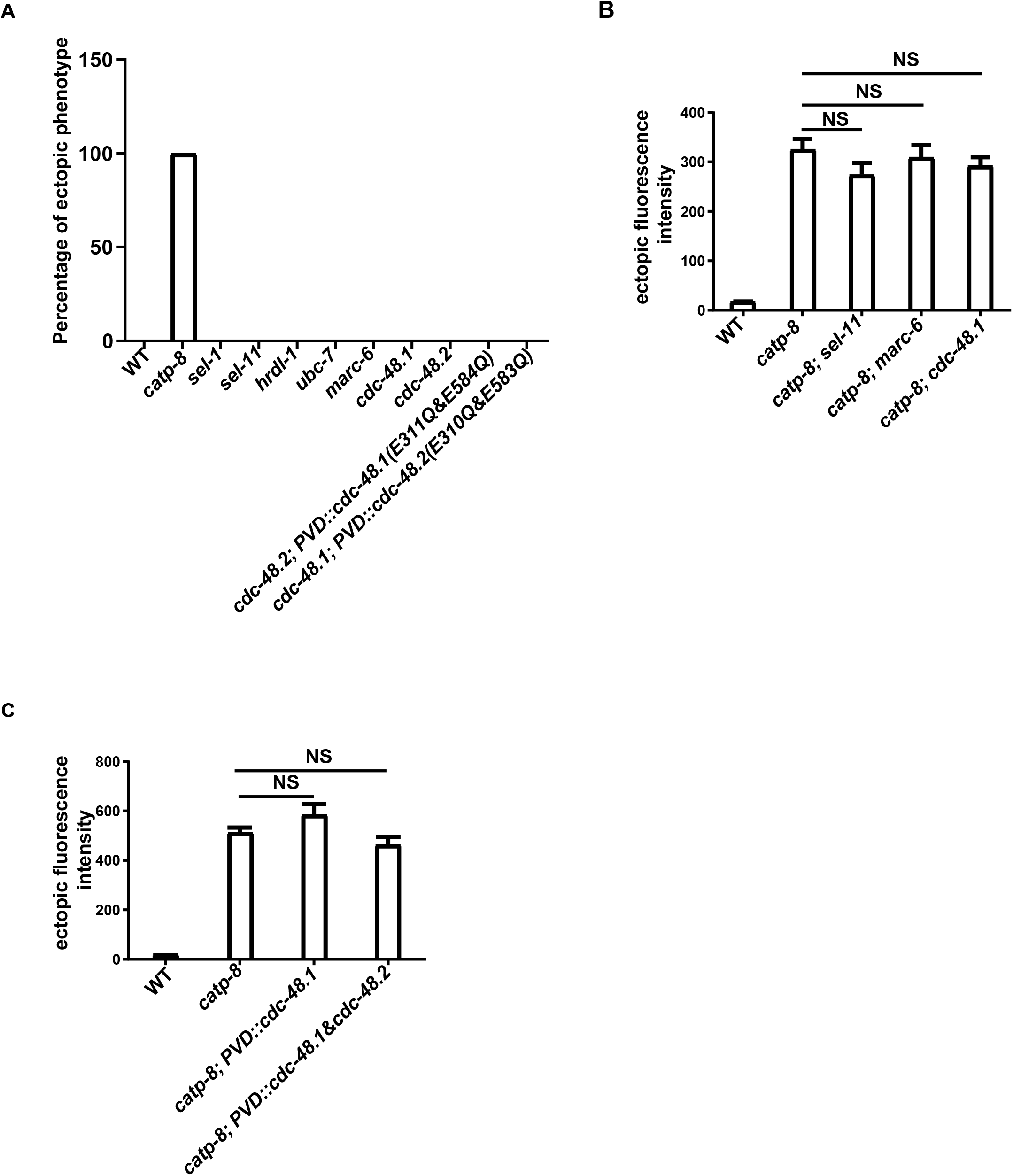
CATP-8 clears mistargeted TOMM-20 from the ER independent of ERAD pathway. **(A)** Quantification of the percentage of TOMM-20 mis-localization phenotype in wild type, *catp-8, sel-1, sel-11, hrdl-1, ubc-7, marc-6, cdc-48.1, cdc-48.2, cdc-48.2;* PVD::CDC-48.1(E311Q&E584Q), and *cdc-48.1;* PVD::CDC-48.2(E310Q&E583Q) mutants. n>=25 for each genotype. **(B)** Quantification of ectopic TOMM-20(1-54AA)::GFP intensity in PVD dendrites in wild type, *catp-8, catp-8; sel-11, catp-8; marc-6, catp-8; cdc-48.1* mutants. Data are shown as mean±SEM. One way ANOVA with Tukey correction. NS means not significant. n>=20 for each genotype. **(C)** Quantification of ectopic TOMM-20(1-54AA)::GFP intensity in PVD dendrites in wild type, *catp-8, catp-8; PVD::cdc-48.1*, and *catp-8; PVD::cdc-48.1&cdc-48.2* strains. Data are shown as mean±SEM. One way ANOVA with Tukey correction. NS means not significant. n>=14 for each genotype.

**Figure S5.**
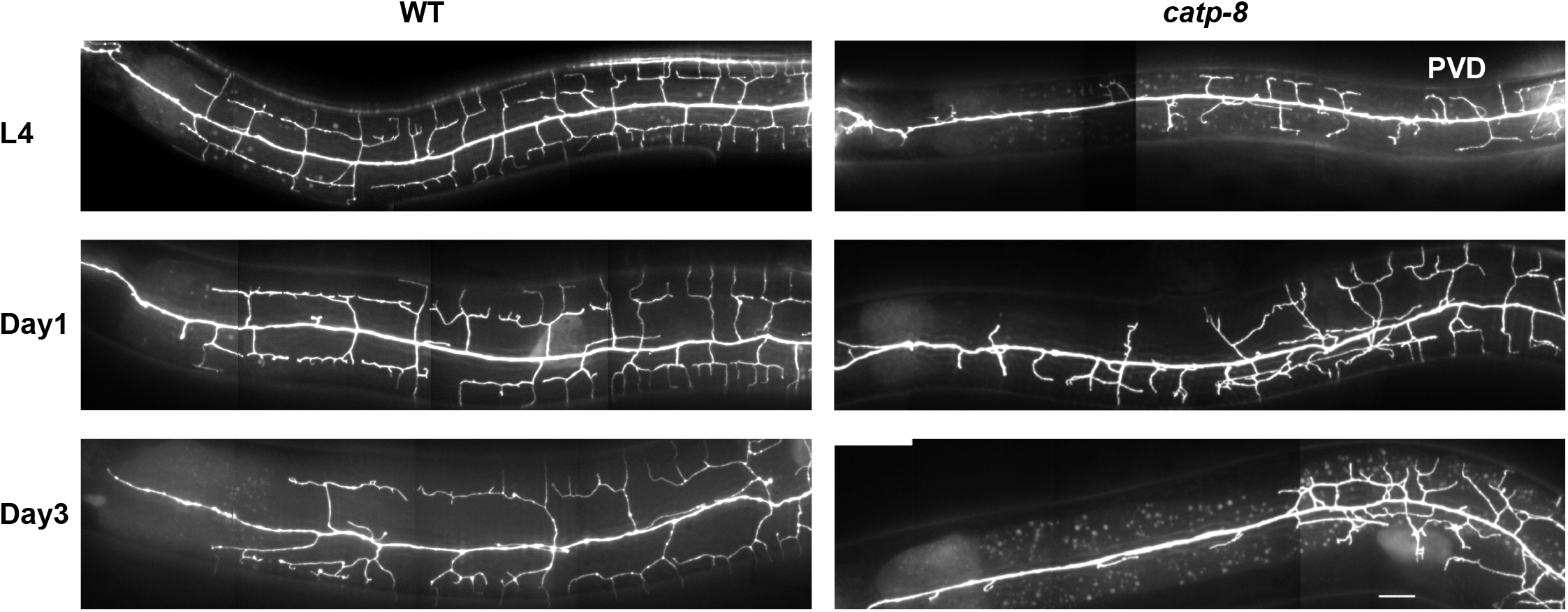
*catp-8* mutant causes developmental but not degenerative dendrite defects. Representative images showing PVD anterior dendrite morphology in wild type and *catp-8* mutant at L4, Day1, and Day3 stage. Scale bar: 10μm.

**Figure S6.**
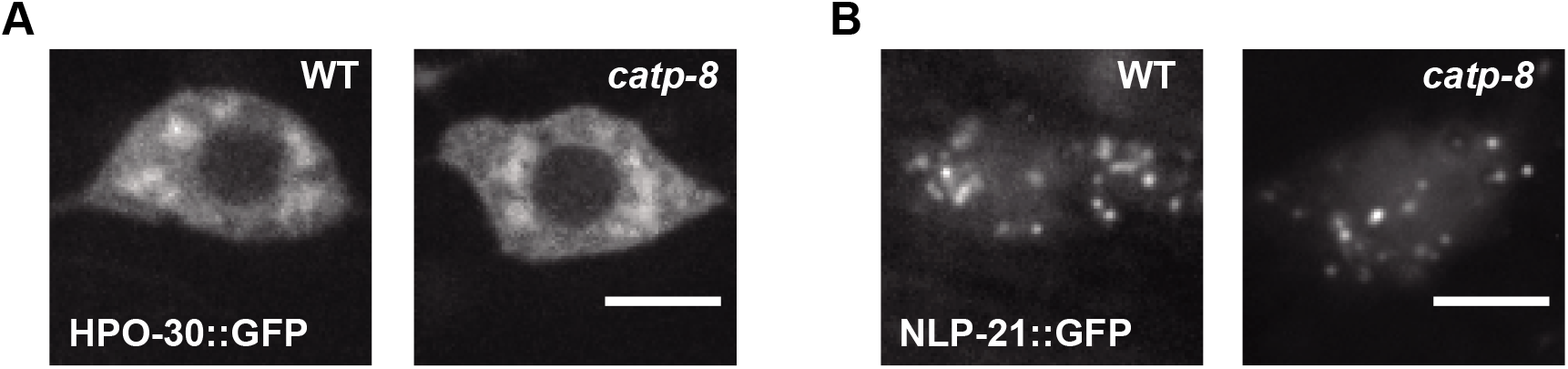
CATP-8 does not affect HPO-30 and NLP-21 expression level in PVD. **(A)** Representative confocal images showing HPO-30::GFP pattern in PVD cell soma in wild type and *catp-8* mutant. Scale bar: 5 μm. **(B)** Representative confocal image showing NLP-21::GFP pattern in PVD cell soma in wild type and *catp-8* mutant. Scale bar: 5 μm.

