## Supplementary tabel 1 for "An endoplasmic reticulum (ER) ATPase safeguards ER identity by removing ectopically localized mitochondrial proteins"

Supplementary Table 1. Strains Used in This Study

| Strain name | Genotype | Source | Notes |
| --- | --- | --- | --- |
| TV51447 | *wyIs50054*  [pXM289(20ng/ul),pOL036(5ng/ul), P*ord-1*::*gfp*(30ng/ul)] | Integration |  |
| TV51183 | *catp-8*(*wy50219*); *wyIs50054* | EMS screening | Q84X |
| TV51777 | *catp-8*(*wy50230*); *wyIs50054* | EMS screening | G1002E |
| TV51725 | *catp-8*(*wy50648*); *wyIs50054* | EMS screening | W71X |
| TV51617 | *catp-8*(*wy50219*);*wyIs50054*; *wyEx50746*[WRM064dE02(50ng/ul), P*ord-1*::*rfp*(50ng/ul)] | Injection |  |
| TV51611 | *wyIs50062*  [pXM289(20ng/ul),pXM190(5ng/ul), P*ord-1*::*gfp*(30ng/ul)] | Integration |  |
| TV51130 | *catp-8*(*wy50219*);*wyIs50062* | Crossing |  |
| TV51618 | *wyEx50736*[pYS36(5ng/ul),pXM190(5ng/ul),P*ord-1*::*gfp*(30ng/ul)] | injection |  |
| TV51263 | *catp-8*(*wy50219*);*wyEx50736* | Crossing |  |
| TV51620 | *wyEx50694*[pXM534(5ng/ul),P*ord-1*::*rfp*(60ng/ul)] | Injection |  |
| TV51066 | *catp-8*(*wy50219*)*;wyEx50694* | Crossing |  |
| TV51834 | *wyEx50844*[pXM532(0.3ng/ul),pXM190(5ng/ul),P*ord-1*::*rfp*(70ng/ul)] | Injection |  |
| TV51835 | *catp-8*(*wy50219*); *wyEx50844* | Crossing |  |
| TV51836 | *wyEx50845*[pXM250(2ng/ul),pXM190(3ng/ul),P*ord-1*::*rfp*(70ng/ul) | Injection |  |
| TV51840 | *catp-8*(*wy50219*); *wyEx50845* | Crossing |  |
| TV51837 | *wySi50064*[pQQ50(50ng/ul)] | Injection |  |
| TV51255 | *catp-8*(*wy50219*);*wySi50064* | Crossing |  |
| TV51828 | *wyIs50077*[pYS103(5ng/ul),P*ord-1*::*rfp*(50ng/ul)];*wyEx50842*[pQQ111(3ng/ul), P*ord-1*::*gfp*(60ng/ul)] | Injection |  |
| TV51829 | *catp-8*(*wy50219*); *wyIs50077*;*wyEx50842* | Crossing |  |
| TV51621 | *wyEx50749*[pQQ54(10ng/ul),P*ord-1*:*:rfp*(60ng/ul)] | Injection |  |
| TV51622 | *catp-8*(*wy50219*);*wyIs50054;wyEx50692*[pQQ27(20ng/ul),P*ord-1*::*rfp*(60ng/ul)] | Injection |  |
| TV51623 | *catp-8*(*wy50219*);*wyIs50054*;*wyEx50758*[pQQ64(20ng/ul),P*ord-1*::*rfp*(60ng/ul)]#1 | Injection |  |
| TV51838 | *catp-8*(*wy50219*);*wyIs50054*;*wyEx50758*[pQQ64(20ng/ul),P*ord-1*::*rfp*(60ng/ul)]#2 | Injection |  |
| TV51624 | *wyEx50806*[pQQ154(15ng/ul),pXM190(5ng/ul),P*ord-1*::*rfp*(40ng/ul)] | Injection |  |
| TV51625 | *wyIs50077*;*wyEx50807*[pQQ134(10ng/ul),P*ord-1*::*gfp*(60ng/ul)] | Injection |  |
| TV51626 | *wyEx50808*[PQQ131(20ng/ul),P*ord-1*::*gfp*(60ng/ul)] | Injection |  |
| TV51627 | *catp-8*(*wy50219*);*wyEx50808* | Crossing |  |
| TV51630 | *catp-8*(*wy50219*);*wyIs50054*;*wyEx50740*[PQQ88a(50ng/ul),P*ord-1*::*rfp*(60ng/ul)] | Injection |  |
| TV51830 | *wyEx50843*[pQQ186(0.1ng/ul),P*ord-1*::*gfp*(60ng/ul)];*wyEx50846*[pXM190(4ng/ul),P*ord-1*::*rfp*(70ng/ul)] | Crossing |  |
| TV51839 | *catp-8*(*wy50219*); *wyEx50843*[pQQ186(0.1ng/ul),P*ord-1*::*gfp*(60ng/ul)];*wyEx50846*[pXM190(4ng/ul),P*ord-1*::*rfp*(70ng/ul)] | Crossing |  |
| TV51831 | *catp-8*(*wy50219*); *wyEx50843*[pQQ186(0.1ng/ul),P*ord-1*::*gfp*(60ng/ul)];*wyEx50740*[pQQ88a(50ng/ul),P*ord-1*::*rfp*(60ng/ul)] | Crossing |  |
| TV51632 | *catp-8*(*wy50219*);*wyEx50809*[PQQ83(1ng/ul);*wyEx50740*[pQQ88a(50ng/ul),P*ord-1*::*rfp*(60ng/ul)] | Crossing |  |
| TV51633 | *sel-1*(*e1948*);*wyIs50054* | Crossing |  |
| TV51258 | *nDf59*;*wyIs50054* | Crossing |  |
| TV51309 | *hrdl-1*(*gk28*);*wyIs50054* | Crossing |  |
| TV51304 | *ubc-7*(*gk5037*);*wyIs50054* | Crossing |  |
| TV51634 | *marc-6*(*wy50647*);*wyIs50054* | Crossing | Deletion |
| TV51301 | *cdc-48.1*(*tm544*);*wyIs50054* | Crossing |  |
| TV51635 | *cdc-48.2*(*tm659*);*wyIs50054* | Crossing |  |
| TV51636 | *cdc-48.2*(*tm659*);*wyIs50054*;*wyEx50810*[PQQ148(20ng/ul),P*ord-1*:*:rfp*(60ng/ul)] | Injection |  |
| TV51637 | *cdc-48.1*(*tm544*);*wyIs50054*;*wyEx50811*[PQQ152(20ng/ul),P*ord-1*::*rfp*(60ng/ul)] | Injection |  |
| TV51250 | *catp-8*(*wy50219*); *nDf59*;*wyIs50054* | Crossing |  |
| TV51638 | *catp-8*(*wy50219*);*marc-6*(*wy50647*);*wyIs50054* | Crossing |  |
| TV51302 | *catp-8*(*wy50219*);*cdc-48.1*(*tm544*);*wyIs50054* | Crossing |  |
| TV51639 | *catp-8*(*wy50219*);*wyIs50054*; *wyEx50812*[PQQ139(20ng/ul),P*ord-1*::*rfp*(60ng/ul)] | Injection |  |
| TV51640 | *catp-8*(*wy50219*);*wyIs50054*;*wyEx50813*[PQQ139(20ng/ul),PQQ141(20ng/ul),P*ord-1*::*rfp*(60ng/ul)] | Injection |  |
| TV51615 | *wyIs50097*  [pXM477(5ng/ul),pOL036(10ng/ul), P*ord-1*::*rfp*(50ng/ul)] | Integration |  |
| TV51296 | *catp-8*(*wy50219*);*wyIs50097* | Crossing |  |
| TV51832 | *catp-8*(*wy50219*);*fis-1*(*tm1867*);*wyIs50097* | Crossing |  |
| TV51833 | *catp-8*(*wy50219*);*fis-1*(*tm1867*);*mff-2*(*tm3041*);*wyIs50097* | Crossing |  |
| TV51297 | *catp-8*(*wy50219*);*drp-1*(*tm1108*);*wyIs50097* | Crossing |  |
| TV15911 | *wyIs592*  [pOL020(20ng/ul),P*ord-1*::*rfp*(50ng/ul)] | Integration |  |
| TV51641 | *catp-8*(*wy50219*);*wyIs592* | Crossing |  |
| TV51242 | *catp-8*(*wy50219*);*wyIs592*;*wyEx50735*[pXM100(20ng/ul), pOL036(20ng/ul)] | Injection |  |
| TV20130 | *wyIs740*  [PVD::*dma-1* ECD::TM::GFP::Cyto(10ng/ul), P*ord-1*::*gfp*] | Integration |  |
| TV51234 | *catp-8*(*wy50219*);*wyIs740* | Crossing |  |
| TV51246 | *catp-8*(*wy50219*);*wyIs740*;*drp-1*(*tm1108*) | Crossing |  |
| NK1531 | *unc-119*(*ed4*), *qyIs366*[PVD::*hpo-30*::*gfp,unc-119*(+)] | Integration |  |
| TV51662 | *catp-8*(*wy50721*);*qyIs366* | Cas9 editing | Deletion |
| TV22696 | *wyIs842*  [PVD::*nlp-21*::Venus (43ng/ul), P*ord-1*::*rfp*] | Integration |  |
| TV51259 | *catp-8*(*wy50219*);*wyIs842* | Crossing |  |
