## Supplementary tabel 2 for "An endoplasmic reticulum (ER) ATPase safeguards ER identity by removing ectopically localized mitochondrial proteins"

Supplementary Table 2. Plasmids Used in This Study

| Plasmid Name | Detailed Information |
| --- | --- |
| pOL036 | *ser-2*prom3::*myri-mCherry* |
| pXM289 | pPD95.77_*ser-2*Prom3::*tomm-20*(1-54aa)::*gfp* |
| pXM250 | pPD95.77_*ser-2*Prom3::*fzo-1*::*gfp* |
| pXM190 | pPD95.77_*ser-2*Prom3::*mCherry*::C34B2.10(rER) |
| pYS103 | pPD95.77_P*myo-3*::*gfp*::*miro-1* |
| pXM532 | pPD95.77_*ser-2*Prom3::*gfp*:: *fis-1* |
| pXM477 | pPD95.77_*ser-2*Prom3::*sec-61.A*::*gfp* |
| pOL020 | *ser-2*prom3::*myri-gfp* |
| pYS36 | pPD95.77_*ser-2*Prom3::*gfp*::*miro-1* |
| pXM534 | pPD95.77_*ser-2*Prom3::*tomm-20*::*gfp* |
| pQQ54 | pPD95.77_P*mbf-1*::*gfp* |
| pQQ27 | pPD95.77_*ser-2*Prom3::*catp-8* |
| pQQ64 | pPD95.77_*ser-2*:Prom3:*catp-8*(D483N) |
| pQQ154 | pPD95.77_*ser-2*Prom3::*gfp*::*catp-8* |
| pQQ134 | pPD95.77_P*myo-3*::*mcherry*::*catp-8* |
| pQQ131 | pPD95.77_*ser-2*Prom3::*stc-1*(signal peptide)::YC3.6::KDEL |
| pQQ88a | pPD95.77_P*hsp-16.48*::*catp-8* |
| pQQ83 | pPD95.77_*ser-2*Prom3::*tomm-20*(1-54aa)::pcSTAR |
| pQQ148 | pPD95.77_*ser-2*Prom3::*cdc-48.1*(E311Q&E584Q) |
| pQQ152 | pPD95.77_*ser-2*Prom3::*cdc-48.2*(E310Q&E583Q) |
| pQQ139 | pPD95.77_*ser-2*Prom3::*cdc-48.1* |
| pQQ141 | pPD95.77_*ser-2*Prom3::*cdc-48.2* |
| pXM100 | pPD95.77_*ser-2*Prom3::*dma-1* |
| pQQ111 | pPD95.77_P*myo-3*::*mcherry*:: C34B2.10(rER) |
| pQQ186 | pPD95.77_*ser-2*Prom3::pcSTAR::*fis-1* |
| pQQ50 | pWZ347_*ser-2*Prom3::*tomm-20*::*gfp* |
